# Histone H3 N-terminal recognition by the PHD finger of PHRF1 is required for proper DNA damage response

**DOI:** 10.1101/2024.11.20.623956

**Authors:** Kanishk Jain, Pata-Eting Kougnassoukou Tchara, Amanuel B. Mengistalem, Aidan P. Holland, Christopher N. Bowman, Matthew R. Marunde, Irina K. Popova, Spencer W. Cooke, Krzysztof Krajewski, Michael-Christopher Keogh, Jean-Philippe Lambert, Brian D. Strahl

## Abstract

Plant homeodomain (PHD) fingers are critical effectors of histone post-translational modifications (PTMs), acting as regulators of gene expression and genome integrity, and frequently presenting in human disease. While most PHD fingers recognize unmodified and methylated states of histone H3 lysine 4 (H3K4), the specific functions of many of the over 100 PHD finger-containing proteins in humans remain poorly understood, despite their significant implications in disease processes. In this study, we undertook a comprehensive analysis of one such poorly characterized PHD finger-containing protein, PHRF1. Using biochemical, molecular, and cellular approaches, we show that PHRF1 robustly binds to histone H3, specifically at its N-terminal region. Through RNA-seq and proteomic analyses, we also find that PHRF1 is intricately involved in transcriptional and RNA splicing regulation and plays a significant role in DNA damage response (DDR). Crucially, mutagenesis of proline 221 to leucine (P221L) in the PHD finger of PHRF1 abolishes histone interaction and fails to rescue defective DDR. These findings underscore the importance of PHRF1-H3 interaction in maintaining genome integrity and provide insight into how PHD fingers contribute to chromatin biology.

## INTRODUCTION

Histone proteins – fundamental to the organization and packaging of the genome – are chemically modified by various “writer” and “eraser” enzymes that install or remove histone post-translational modifications (PTMs), respectively (1, 2). These histone PTMs play a central role in chromatin function, largely through their ability to recruit effector or “reader” domain-containing proteins that interact with histones, DNA and other chromatin-associated proteins to regulate gene transcription and other DNA templated functions such as DNA repair (2, 3). The diverse patterns of histone PTMs, referred to as the Histone Code, are recognized by different reader proteins, which modulate chromatin structure and function (2, 3). Dysregulation of the epigenetic machinery, notably the readers and writers, is implicated in a wide range of human diseases, including cancer (4). Despite significant progress, many annotated reader domains have gone uncharacterized, and even less is understood about how tandem readers in chromatin-associated proteins function in a combinatorial manner. Given the link between plant homeodomain (PHD) fingers and diseases such as breast cancer and leukemia (5–10), there is a pressing need to elucidate their histone-binding preferences and roles in chromatin biology.

In an effort to elucidate the role of PHD fingers, we recently examined the histone binding preference(s) of multiple members of the PHD domain family (11). This work identified several PHD-histone interactions that were previously unknown, one of which was the interaction of PHRF1 (PHD and RING Finger protein 1) with the unmodified N-terminus of histone H3. PHRF1 is an E3 ubiquitin ligase that contains a C-terminal SRI (Set2-Rpb1 Interacting) domain known to bind the C-terminal tail of RNA polymerase II (RNAPII) (12–14) (Figure 1A). Several reports suggest that PHRF1 plays significant roles in cancer biology. For example, overexpression of PHRF1 in MCF-7 breast cancer xenografts almost completely abrogates tumor growth (12), highlighting its potential therapeutic relevance. Conversely, overexpression of PHRF1 in lung cancer showed the opposite effect, suggesting context-dependent roles for this enzyme in cancer (15). Additionally, genome-wide association studies have implicated PHRF1 in autoimmune diseases such as systemic lupus erythematosus (16, 17), and recent work has implicated its ubiquitination activity in non-homologous end joining DNA damage repair (18).

**Figure 1.**
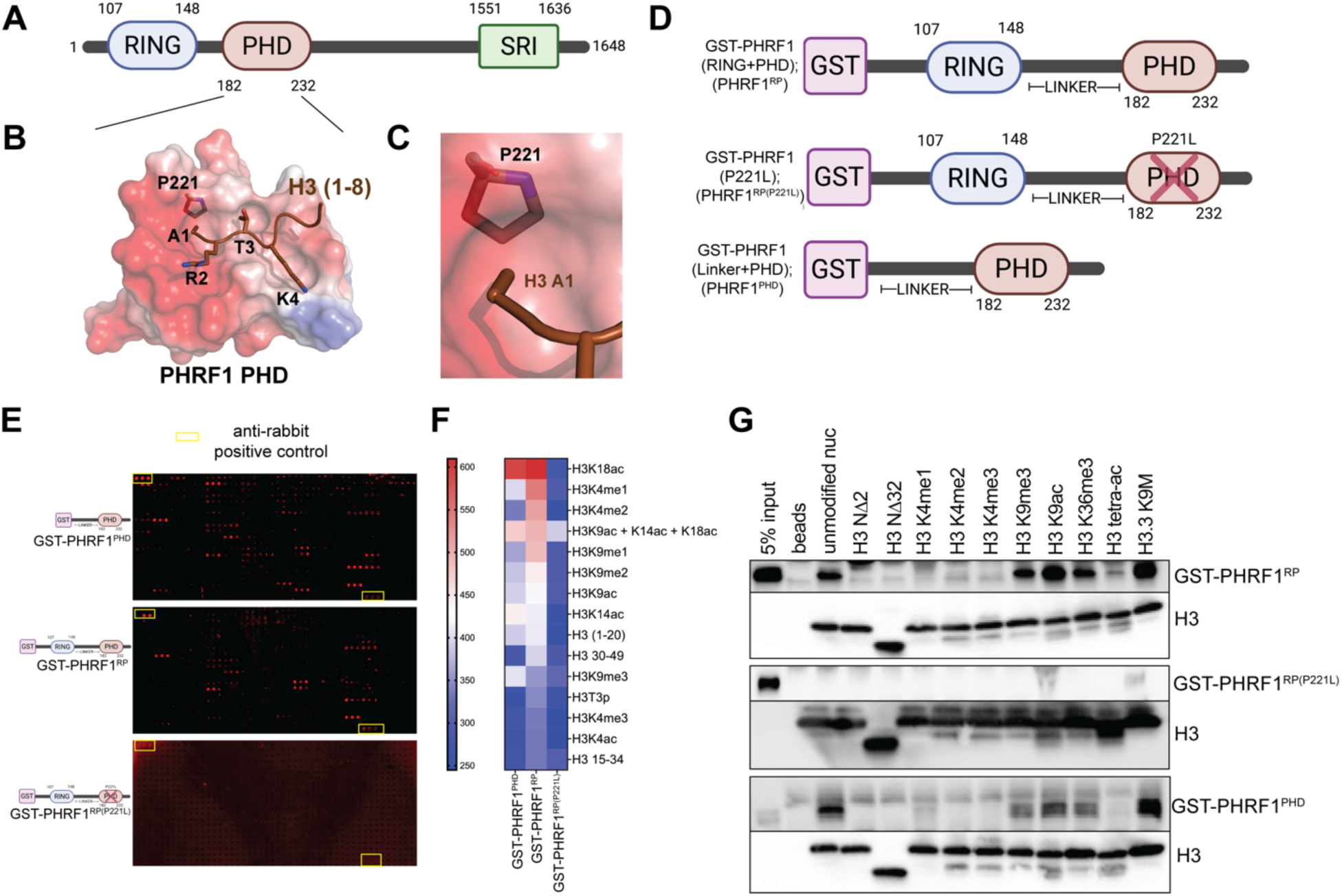
The PHD finger of PHRF1 binds to the N-terminus of histone H3. **A.** Domain schematic of the protein PHRF1, highlighting the RING finger (purple oval), PHD finger (brown oval), and SRI region (green rectangle). **B.** Structural homology model of the PHD region of PHRF1 (shown as a translucent protein surface with electrostatics calculated through APBS in PyMOL) docked to a peptide comprised of the first eight residues of histone H3 (ARTKQTAR), shown as a peptide in brown. P221 of PHRF1 and the first 4 residues of the H3 peptide are also shown as stick representations. This model was generated in Alphafold-Multimer. **C.** A zoomed in region of the PHRF1 PHD:H3 (1–8) structural homology model highlighting a binding pocket formed by PHRF1 P221 for H3 A1 to enter. **D.** A sequence schematic for GST-tagged PHRF1 constructs for *in vitro* experiments. **E.** Biotinylated histone peptide microarray binding images with GST-PHRF1^PHD^ (top), GST-PHRF1^RP^ (middle), and GST-PHRF1^RP(P221L)^ (bottom). Triplicate red dots indicate a positive histone peptide binding event (each peptide is arrayed in triplicate). Regions outlined by yellow boxes indicate anti-rabbit positive controls. **F.** A heatmap comparing binding intensities observed in the peptide microarrays for each of the three GST-PHRF1 constructs to select peptides (indicated on the y-axis). Binding strength is represented on a color gradient from red to blue (stronger to weaker) (n=4). For full peptide microarray data, please see Supplementary File 1. **G.** Representative western blots of biotinylated nucleosomes pulldowns between differentially modified nucleosomes (top) and GST-PHRF1 constructs (right). Anti-GST blots represent PHRF1 signal and anti-H3 blots are shown as loading controls. Key: H3 NΔ2 and NΔ32 are nucleosomes lacking residues 1-2 pr 1-32 on H3, respectively, and H3 tetra-ac = H3 K4acK9acK14acK18ac.

While these discoveries underscore the biological significance of PHRF1, there remains a lack of thorough characterization of this protein’s functions in chromatin biology, particularly in human disease. Given that PHRF1 harbors canonical chromatin-interacting domains such as a PHD finger, it is crucial to understand how interaction of PHRF1 with histones contributes to its biological functions. With this in mind, we sought to further understand the histone binding activity of PHRF1 and to ask if this binding is functionally relevant to the biological roles of PHRF1. Since our understanding of PHRF1 is also very limited, a holistic approach to define its roles in transcription and DNA repair is warranted.

In this report, we comprehensively examine the histone interactions of PHRF1 and further define how this E3 ubiquitin ligase contributes to transcriptional regulation as well as DNA damage response (DDR). We provide evidence that H3 reading by PHRF1 plays a critical role in DDR, highlighting the importance of PHRF1’s PHD finger to chromatin-based events. As PHRF1 is overexpressed in multiple cancers, our findings suggest the PHD-histone reading activity of PHRF1 will be important to its contribution to cancer.

## MATERIALS AND METHODS

### Antibodies

GST (in-house generated, 1:1000), H3 (in-house generated, 1:5000), PHRF1 (in-house generated, 1:500), GAPDH (Cell Signaling Technology (CST) 5104, 1:5000), γH2A.X (CST 80312, 1:200), 53BP1 (CST 4937, 1:400), RNAPII (Active Motif 39497, 1:500), SRSF1 (ThermoFisher 32-4600, 1:2000), PARP1 (CST 9542, for IB: 1:1000, for IF: 1:250).

### Structural homology modeling

The amino acid sequence for the PHD region of human PHRF1 (Uniprot ID: Q9P1Y6; residues 182-233) and the N-terminus of human histone H3.1 (Uniprot ID: P6843; residues 1-8) were used as input queries in Alphafold2 (Alphafold-Multimer) (19). Once models were generated, the best model was visualized in PyMOL (Schrodinger, LLC). For visualization purposes, PHRF1’s PHD was represented as a protein’s globular surface, with electrostatic potentials calculated via the APBS plugin in PyMOL.

### Generating GST-PHRF1 constructs

Constructs shown in Figure 1D were all derived from the GST-PHRF1^RP^ (containing residues 107-232 of PHRF1) pGEX6T plasmid initially reported in an earlier study (11). Specifically, the RING domain (residues 107-148) was deleted to generate GST-PHRF1 Linker + PHD (GST-PHRF1^PHD^) and P221 was mutated to a leucine residue in the GST-PHRF1^RP^ construct to create GST-PHRF1^RP(P221L)^ using a QuikChange II Site-Directed Mutagenesis Kit from Agilent Technologies. All constructs were purified and quantified identically, as described previously (11).

### Histone peptide microarrays

Histone peptide microarrays with GST-tagged PHRF1 constructs from Figure 1D were performed and quantified as previously reported (11). Experiments were carried out in quadruplicate. See Supplementary File 1 for full results.

### Histone peptide fluorescence polarization

Fluorescence polarization with GST-PHRF1 constructs and a C-terminally fluorescently labeled H3 peptide (ARTKQTARKSTGGKAPRKQL-K(5-FAM)-NH2; H3 (1–20)-FAM) was carried out similarly to a previously described protocol (20). Briefly, GST-PHRF1^RP^, GST-PHRF1^PHD^, and GST-PHRF1^RP(P221L)^ proteins were titrated from 50 to 0.0061 μM and 0 μM and incubated with 50 nM H3 (1–2)-FAM 20 mM Tris (pH 8.0), 250 mM NaCl, 1 mM DTT, and 0.05% NP-40 for 20 minutes in 40 μL reactions in black, flat-bottomed 384-well plates (Costar). Fluorescence polarization was measured on CLARIOstar Plus plate reader (BMG Labtech) using an excitation filter of 482 nm and emission filter of 515-530 nm. In order to determine dissociation constants (KD), data were graphed and fit to a one-site binding model in GraphPad Prism 10.0.

### dCypher nucleosome alphascreen binding assays

dCypher binding assays were performed using GST-PHRF1^RP^ (3nM) and differentially modified biotinylated nucleosomes (10 nM; EpiCypher Inc.). Binding was detected using AlphaScreen technology (Revvity) and all reactions were carried out in duplicate and performed as in Jain *et al*. (21).

### Nucleosome pulldown assays

Nucleosome pulldown assays were performed using a binding buffer (NBB) composed of 50 mM Tris-HCl pH 7.5, 150 mM NaCl, 0.5% bovine serum albumin (BSA), 0.1% NP-40, and 10% glycerol. For each binding reaction, 2.5 pmol of protein (GST-PHRF1^RP^, GST-PHRF1^PHD^, or GST-PHRF1^RP(P221L)^) were diluted in 22.5 µL of NBB and mixed with 12.5 pmol of differentially modified biotinylated nucleosomes (EpiCypher Inc.); negative controls without nucleosomes were included. The mixtures were rotated at 4°C overnight. Streptavidin magnetic beads (1 µL per reaction; Invitrogen Dynabeads M-280 Streptavidin) were washed twice with NBB and resuspended in 7.5 µL of NBB per reaction before being added to the samples and rotated for 1 hour. Beads were then washed three times with 200 µL of NBB, rotating for 5 minutes at 4°C during each wash, and finally resuspended in 15 µL of 1× SDS loading dye for immunoblot analysis, using an anti-GST antibody to detect PHRF1 and an anti-H3 antibody for loading/presence of nucleosomes.

### Biolayer Interferometry (BLI)

Biolayer interferometry (BLI) was carried out using GST-PHRF1^RP^ and GST-PHRF1^RP(P221L)^ and biotinylated unmodified nucleosomes (25 nM) immobilized on streptavidin biosensors (Sartorius). All samples were prepared in 20 mM MOPS (pH 7.0), 150 mM KCl, 1 mM DTT, and 0.2 mg/mL BSA as reported previously (22). Streptavidin biosensors were hydrated in the BLI buffer for 30 minutes prior to data collection. Experiments were performed at 37°C in black, 96-well plates (Greiner Bio-One), while shaking at 1,000 rpm. Importantly, data were collected at 10 Hz, with an experimental scheme as follows: 10 minutes of temperature pre-equilibration, 180s of buffer equilibration, 300s of biotinylated nucleosome loading, 120s of baseline equilibration, 300s of GST-protein association with nucleosomes, 300s of dissociation. GST-PHRF1^RP^ concentrations used were 20, 10, 5, 2.5, 1.25 and 0 µM. GST-PHRF1^RP(P221L)^ concentrations used were 25, 12.5, 6.25, 3.125, 1.625, and 0 µM. Data were collected on a ForteBio Octet Red834 BLI instrument and were analyzed using the Octet Analysis Software in order to determine KD values. Data was background subtracted using the 0 µM runs.

### Cell lines and culture conditions

The following human cell lines were obtained from the American Type Culture Collection (ATCC, Manassas, VA, USA): MCF-7 (ATCC HTB-22), BT-20 (ATCC HTB-19), T47-D (ATCC HTB-133), NCI-H1299 (ATCC CRL-5803), NCI-H1650 (ATCC CRL-5883), A549 (ATCC CCL-185), HCT116 (ATCC CCL-247), HEK293T (ATCC CRL-11268), NCI-N87 (ATCC CRL-5822), U87-MG (ATCC HTB-14), SKOV3 (ATCC HTB-77), and HeLa (ATCC CCL-2). The SNU1040 cell line was obtained from Dr. Tom Smithgall (University of Pittsburgh, Pittsburgh, PA) and the HKC cell line was obtained from Dr. Lorraine Racusen (Johns Hopkins Hospital, Baltimore, MD). HEK293T, HKC, MCF-7, HeLa, BT-20, and U87-MG cells were cultured in Dulbecco’s Modified Eagle’s Medium (DMEM), with MCF-7 cells additionally supplemented with 0.01 mg/mL human recombinant insulin. T47-D cells were grown in RPMI-1640 medium supplemented with 0.2 units/mL human recombinant insulin. NCI-H1299, NCI-H1650, NCI-N87, and SNU1040 cells were cultured in Roswell Park Memorial Institute (RPMI)-1640 medium. A549 cells were grown in F-12K. HCT116 and SKOV3 cells were maintained in McCoy’s 5A medium. All cell lines were also supplemented with 10% fetal bovine serum (FBS), 100 U/mL penicillin, and 100 µg/mL streptomycin, and were incubated at 37 °C in a humidified atmosphere containing 5% CO2.

### Whole cell lysate preparation

For preparation of whole-cell lysates (WCLs) from human cells, cells were resuspended in RIPA buffer containing 25 mM HEPES (pH 8.0), 150 mM NaCl, 1% NP-40, 0.1% SDS, and 0.5% sodium deoxycholate, supplemented with freshly added protease inhibitors [phenylmethylsulfonyl fluoride (PMSF), pepstatin, leupeptin, and aprotinin]. Protease inhibitors were added from stock solutions at 1,000× concentration for aprotinin, leupeptin, and pepstatin, and 100× for PMSF (0.1 M) homogenized by pipetting up and down. Universal Nuclease (250 U per sample; Pierce) was added, and the lysates were rotated at 4 °C for 1.5 hours. Lysates were then centrifuged at 13,000 rpm for 2 minutes at 4 °C, and the supernatants (WCLs) were collected. Protein concentrations were determined using the detergent-compatible Bradford assay (ThermoFisher). WCLs were further analyzed by immunoblotting/western blot analysis.

### PHRF1 co-immunoprecipitation

For immunoprecipitation of PHRF1, HeLa cells were lysed in lysis buffer containing 50 mM HEPES (pH 8.0), 150 mM NaCl, and 1% NP-40, supplemented with freshly added protease inhibitors (PMSF, pepstatin, leupeptin, and aprotinin). Universal Nuclease (1000U per sample) was added, and the lysates were incubated with gentle rotation at 4 °C for 1 hour. After centrifugation at 13,000 rpm for 5 minutes at 4 °C, the supernatants (whole-cell lysates, WCLs) were collected. WCLs were pre-cleared by incubating with 40 μL of equilibrated Protein G agarose beads (ThermoFisher) for 30 minutes at 4 °C, followed by centrifugation to remove the beads. The pre-cleared WCLs were then split equally into two samples: one for immunoprecipitation with anti-PHRF1 antibody and the other with control IgG. Separately, 5 μg of anti-PHRF1 antibody (in-house) or normal rabbit IgG (CST; 2729) was incubated with 35 μL of Protein G Dynabeads (ThermoFisher) in lysis buffer containing 1% BSA for 2 hours at 4 °C to immobilize the antibodies. The respective WCLs were added to the antibody-conjugated beads and incubated overnight at 4 °C with rotation. Beads were washed three times with 1 mL of cold PBS, and bound proteins were eluted by resuspending the beads in 100 μL of 1× SDS loading dye. Samples were then prepared for SDS-PAGE and Western blot analysis.

### Cell line generation for BioID mass spectrometry

For BioID experiments, Flp-In T-REx HeLa cells were used and a construct encoding miniTurbo-tagged PHRF1 was generated via Gateway cloning into pDEST 3′ miniTurbo-3XFlag pcDNA5-FRT-TO as per (23). MiniTurbo-3XFlag - and miniTurbo-3XFlag -GFP expressing cells were used as negative controls and processed in parallel. BioID stable cell lines were selectively grown in the presence of 200 μg/mL hygromycin until 80% confluent, when expression was induced via 1 μg/mL tetracycline for 23 h. Then, the cells were treated with 50 μM biotin for 1 h and harvested and stored at -80°C until streptavidin purifications.

### Proximity-dependent biotinylation (BioID)

The BioID protocol was adapted from (23), with slight modifications. Cells from two 15 cm plates were pelleted, frozen, and thawed in 1.5 mL ice cold radioimmunoprecipitation buffer (RIPA; 50 mM Tris-HCl (pH 7.5), 150 mM NaCl, 1% NP-40, 1 mM ethylenediaminetetraacetic acid, 1 mM ethylene glycol tetraacetic acid, 0.1% sodium dodecyl sulfate, and 0.5% sodium deoxycholate). Phenylmethylsulfonyl fluoride (1 mM), DTT (1 mM), and Sigma-Aldrich protease inhibitor cocktail (1:500) were added immediately before use. The samples were sonicated in 30 s bursts with 2 s pauses every 10 s at 35% amplitude using a Q125 sonicator (QSONICA). Then, turbonuclease (100 units, Sigma-Aldrich, T4332) was added and the lysates were rotated at 4°C for 1 h. For each sample, 60 μL of Streptavidin Sepharose High Performance Affinity Chromatography Medium (Cytiva, Cat 17-5113-01) was prewashed three times with 1 mL of RIPA buffer, by pelleting the beads with gentle centrifugation and aspirating the supernatant before adding the next wash. Biotinylated proteins were captured on pre-washed streptavidin beads for 3 h at 4°C with rotation. The beads were gently pelleted and then washed twice with 1 mL of RIPA buffer and three times with 1 mL of 50 mM ammonium bicarbonate (pH 8.0). Following the final wash, the beads were pelleted and any excess liquid was aspirated. Beads were resuspended in 100 μL of 50 mM ammonium bicarbonate, and 1 μg of trypsin solution was added. The samples were rotated overnight at 37°C and then an additional 1 μg of trypsin was added, followed by an additional 2–4 h of incubation. The beads were pelleted and the supernatant was transferred to a fresh tube. The beads were rinsed twice with 100 μL of high-performance liquid chromatography-grade acetonitrile and the wash fractions were combined with the supernatant. The peptide solution was acidified with 50% formic acid to a final concentration of 2% and the samples were dried in a SpeedVac. Samples were acidified with formic acid to a final concentration of 2% and desalted using homemade C18 Stage Tips as previously described (24). Tryptic peptides were stored at -80°C until MS analysis.

### Experimental design and statistical rationale for MS experiments

For each analysis, at least two biological replicates of each tagged proteins were processed independently, with negative controls included in each batch of processed samples. The order of sample acquisition on the LC-MS/MS system was randomized. Statistical scoring was performed against the negative controls using LFQ-Analyst using default settings (25).

### Data dependent acquisition MS

MS analyses were performed at the Proteomics Platform of the Quebec Genomics Center. Peptide samples were separated by online reversed-phase nanoscale capillary liquid chromatography and analyzed by electrospray MS/MS. The experiments were performed with a Dionex UltiMate 3000 RSLCnano chromatography system (Thermo Fisher Scientific) connected to an Orbitrap Fusion mass spectrometer (Thermo Fisher Scientific) equipped with a nanoelectrospray ion source. Peptides were trapped at 20 μL/min in loading solvent (2% acetonitrile, 0.05% TFA) on an Acclaim 5μm PepMap 300 μ-Precolumns Cartridge Column (Thermo Fisher Scientific) for 5 min. Then, the precolumn was switched online with a laboratory-made 50 cm × 75 μm internal diameter separation column packed with ReproSil-Pur C18-AQ 3-μm resin (Dr. Maisch HPLC) and the peptides were eluted with a linear gradient of 5–40% solvent B (A: 0,1% formic acid, B: 80% acetonitrile, 0.1% formic acid) over 90 min at 300 nL/min. Mass spectra were acquired in data-dependent acquisition mode using Thermo XCalibur software version 3.0.63. Full scan mass spectra (350–1,800 *m/z*) were acquired in the Orbitrap using an AGC target of 4e5, a maximum injection time of 50 ms, and a resolution of 120,000. Internal calibration using lock mass on the *m/z* 445.12003 siloxane ion was used. Each MS scan was followed by acquisition of the fragmentation spectra of the most intense ions for a total cycle time of 3 s (top speed mode). The selected ions were isolated using the quadrupole analyzer in a window of 1.6 *m/z* and fragmented by higher energy collision-induced dissociation at 35% collision energy. The resulting fragments were detected by the linear ion trap in rapid scan rate with an AGC target of 1e4 and a maximum injection time of 50 ms. Dynamic exclusion of previously fragmented peptides was set for a period of 20 sec and a tolerance of 10 ppm.

### Protein identification and enrichment quantification

MaxQuant (version 2.0.3.0; http://www.maxquant.org/) was used to identify and quantify proteins in the dataset, using default parameters with a few exceptions. Peptides were identified from MS/MS spectra using UniProt Human Proteome UP000005640_9606 (UniProt release 2023_03, containing 20,586 proteins) as a reference database. Fractions from the same sample were set as individual experiments in the same parameter group. Replicates were processed independently. Methionine oxidation and N-terminal acetylation were set as variable modifications. Trypsin was selected as the protease, with up to two missed cleavages allowed. The peptide length range was set from 7–25, with a maximum mass of 4600 Da. The false discovery rate was set to 0.01 for peptides, proteins, and sites. The analysis was performed using label-free quantification with MaxLFQ implemented in MaxQuant (26). The “proteinGroups” .txt files produced by MaxQuant were used as input files to analyze and visualize the data in volcano plots with LFQ-Analyst using default settings (25). Detailed MS data are provided in Supplementary File 2.

### CRISPR-Cas9-directed knockout of PHRF1

A predesigned synthetic guide RNA (sgRNA) against PHRF1 was purchased from Integrated DNA Technologies (IDT) with a sequence of 5’ GGAGAACACCAAAGCGAGCG 3’. CRISPR-Cas9 ribonucleoprotein (RNP) complexes were prepared fresh by mixing 50 pmol Cas9 protein (61µM) with 100 pmol sgRNA, bringing up to a volume of 4 µL in 1X phosphate buffered saline (PBS), incubating at room temperature for 20 minutes. 5×10^5^ cells were trypsinized, washed in 1X PBS, and resuspended in 20 µL of supplemented SE Nucleofector Solution as per the manufacturer’s guidelines for HeLa and HCT116 cells (Lonza Bioscience). The entire RNP complex solution and 1 µL 100 µM Alt-R Cas9 electroporation enhancer (IDT) were added to each 20 µL cell aliquot before being transferred to a nucleocuvette (Lonza Bioscience). The following electroporation protocols were used on a Lonza 4D-Nucleofector System: CN114 for HeLa and EN113 for HCT116. Cells were then transferred to the appropriate pre-warmed media in a well from a 12-well tissue culture plate and allowed to recover at 37°C. Once the cells were confluent, they were diluted in 15cm tissue culture dishes and resultant colonies were picked/screened for PHRF1 expression by immunoblotting (data not shown). Clones showing lack of PHRF1 protein relative to control/untreated cells were expanded for use as ΔPHRF1 cells.

### RNA extraction, sequencing, analysis, and rMATS

Total RNA was extracted from 1×10^6^ control and ΔPHRF1 cells in biological triplicate (for both HeLa and HCT116) using TRIzol reagent (Invitrogen), following the manufacturer’s protocol. Purity and concentration were initially assessed using a NanoDrop spectrophotometer. cDNA library preparation and next-generation sequencing were performed by Novogene. Specifically, 150 nt paired-end sequencing was performed with read depth of 80 million reads per replicate and 20 million reads per replicate for HeLa and HCT116 cells, respectively. Raw sequencing reads in FASTQ format were aligned to the human reference genome (GRCh38) using STAR aligner (version 2.7.3a) (27). Alignment and transcript quantification were performed using Partek Flow software (version 10.0; Partek Incorporated, St. Louis, MO, USA). Transcript quantification utilized the Partek E/M algorithm within Partek Flow. Differential gene expression analysis was carried out using DESeq2 (version 1.26.0) (28) within the Partek Flow environment. Lists of differentially expressed genes for HeLa and HCT116 cells (ΔPHRF1 versus control) can be found in Supplementary Files 4 and 8, respectively. For HeLa RNA-seq, alternative splicing events were determined using replicate Multivariate Analysis of Transcript Splicing (rMATS, version 4.0.2) (29). For full list of differential alternative splicing events, please see Supplementary File 6.

### GO-term/Pathway analyses

Gene ontology (GO)-term and pathway analyses for proteomic and genomic data were performed using a variety of tools. For BioID GO-term analysis, proteins that were significantly enriched with PHRF1 (p-value <0.05, log2(fold-change) >1; Supplementary File 2) were entered into DAVID Bioinformatic Resources (30). Specifically, the following databases were searched: UP_KW_Biological_Process, GOTERM_CC_DIRECT, GOTERM_MF_DIRECT, and GOTERM_BP_DIRECT (see Supplementary File 3 for full list). Ingenuity Pathway Analysis (IPA; Qiagen, Inc.) was used for pathway analysis of RNAseq data from HeLa and HCT116 cells. Specifically, for the HeLa RNAseq data, the input differentially expressed genes (DEGs) list was generated after applying an FDR cutoff of 0.0001, and for HCT116 cells an FDR cutoff of 0.05. Pathway enrichment was determined using the Ingenuity Knowledge Base within the IPA software (see Supplementary Files 5 and 9 for full lists). Finally, after rMATS analysis was performed on aligned reads from the HeLa RNAseq experiment, a filtered list (FDR <0.05 and ΔΨ >0.1) was used for functional analysis. The GO Biological Process database on Enrichr (31) was used to analyze significantly enriched biological processes (see Supplementary File 7). For all methods discussed, the enrichment of select terms/pathways was represented in bubble plots generated using RStudio 4.1.1 and the ggplot2 package.

### PHRF1 expression in cancer cells and overall survival in cancers

PHRF1 expression data were obtained from the DepMap database (24Q2 release) (32, 33), which provides gene expression profiles for a wide range of cancer cell lines. Lineages corresponding to individual cell lines were then categorized into broader tissue types. Specifically, each lineage term was assigned to one of the following primary tissue types: Blood/Immune, Digestive, Reproductive, Skin/Soft Tissue, Urinary, Respiratory, Nervous, Endocrine, Musculoskeletal, or Other. For example, the Breast lineage was categorized under Reproductive, while Myeloid and Lymphoid lineages were assigned to the Blood/Immune category. After categorization, expression values corresponding to each lineage were aggregated within their respective tissue type categories. A box-whisker plot was generated in R using the ggplot2 package to visualize the distribution of expression values across the categorized tissue types. Jittered individual data points were overlaid to highlight the variability within each category. Median expression values were calculated and used to sort the tissue types from lowest to highest, with plots subsequently saved as SVG files for inclusion in the final manuscript. Additionally, a Kaplan-Meier (KM) survival plot for overall cancer survival based on PHRF1 expression was generated using GEPIA, which utilizes data from The Cancer Genome Atlas (TCGA) (34) and Genotype-Tissue Expression (GTEx) (35) projects to assess the prognostic value of gene expression in various cancers.

### Generation of stable ΔPHRF1 complementation cell lines

HCT116 ΔPHRF1 cells were infected with lentivirus containing plasmids expression N-terminally 3XFlag-tagged PHRF1 (3XFlag-PHRF1^WT^), PHRF1^P221L^ (3XFlag-PHRF1^P221L^), or a 3XFlag containing empty vector (EV), followed by selection with blasticidin (7.5 µg/mL); HCT116 control cells were also infected with eh EV lentivirus. Selection was conducted alongside HCT116 cells that were not infected with lentivirus containing blasticidin resistance as a gauge for efficient selection. Once selection was confirmed, clones were isolated identically to those for ΔPHRF1 cells earlier, through dilution, picking, and assessing of the PHRF1 protein via immunoblotting compared to the control and ΔPHRF1 cells (data not shown). Clones showing continued lack of PHRF1 protein expression (along with blasticidin resistance) were designated ΔPHRF1+EV and clones with similar PHRF1 protein expression to that seen in control HCT116 cells were designated as either Control+EV, ΔPHRF1+3XFlag-PHRF1 ^WT^ or ΔPHRF1+3XFlag-PHRF1^P221L^.

### Live-cell imaging growth analysis of HCT116 cell lines

Control+EV, ΔPHRF1+EV, ΔPHRF1+3XFlag-PHRF1^WT^, and ΔPHRF1+3XFlag-PHRF1^P221L^ HCT116 cells were harvested at approximately 70% confluence, counted, and seeded into 96-well plates at a density of 1,000 cells per well in 100 µL growth medium (see above), with each cell line seeded in biological triplicates. Immediately after seeding, plates were placed in the IncuCyte S3 Live-Cell Analysis System for live-cell imaging. Images were captured every 4 hours over one week using a 10X objective lens, with four fields of view per well. The IncuCyte S3 software quantified cell confluence as the percentage of area occupied by cells; data from the four fields per well were averaged at each time point and then averaged across triplicates for each cell line. Confluence data were exported to GraphPad Prism 10 for plotting growth curves; for each cell line, data were individually normalized to their confluence at time zero to account for any initial differences.

### Zeocin DNA damage recovery assay, immunofluorescence, and immunoblotting

To induce DNA double-strand breaks (DSBs), cells were first seeded at a density of 10,000 cells per well 96-well µClear plates (Greiner Bio-One) in triplicate and grown to confluence for 2 days. Cells were then treated with zeocin (Invitrogen) under specific conditions. Control and ΔPHRF1 HeLa cells were exposed to 50 µg/mL zeocin for 30 minutes at 37°C, whereas Control, ΔPHRF1, and complemented HCT116 cells (Control+EV, ΔPHRF1+EV, ΔPHRF1+3XFlag-PHRF1^WT^, and ΔPHRF1+3XFlag-PHRF1^P221L^) were treated with 10 µg/mL zeocin for the same duration and temperature. Following treatment, cells were washed three times with pre-warmed phosphate-buffered saline (PBS) to remove residual zeocin. Fresh complete medium was added, and the cells were allowed to recover at 37°C. Samples were fixed in 4% methanol-free formaldehyde (ThermoFisher) for 15 min at room temperature at designated time points during the recovery phase—specifically at untreated, 0, 1, 4, 11, and 26 h for HeLa cells and untreated, 0, 1, 4, 8, and 12 h for HCT116 cells post-washout—for subsequent analyses. Following fixation, the cells were washed three times in 1X PBS and then permeabilized in 0.5% Triton X-100 in 1XPBS for 10 minutes at room temperature. Cells were subsequently washed three times with PBS. Cells were then blocked in 6% BSA in 1XPBS for 30 min at room temperature before incubation in primary antibody for 3 h at room temperature. After washing the cells in PBS again, the cells were incubated in secondary antibodies (anti-rabbit AlexaFlour 488 and anti-mouse AlexaFluor 594) at 1:500 for 1.5 h at room temperature in the dark. Cells were washed and stained with DAPI. Cells were again washed and stored in 1XPBS until imaging. In each cell line discussed, PHRF1 protein expression at each time point was assessed by western blot analysis of whole cell lysates made in RIPA buffer.

### Microscopy and Imaging Analysis

For purposes of quantification, widefield images were acquired using a GE IN CELL Analyzer 2200 with a 40X/0.95 Plan-Apo objective. Images were processed in ImageJ and CellProfiler. For purposes of visualization, confocal images were acquired on a Zeiss 880 confocal microscope using a Plan-Apo 20X/0.8 WD 0.55mm at 4X zoom. Z-stacks of 0.9 µm slices were taken and maximum intensity projections were made in ImageJ for display. Signal intensities and DNA damage foci from widefield microscopy were quantified using CellProfiler (version 4.2.4)(36). First, flat-field correction was applied to all images using CellProfiler and ImageJ. From the corrected images, nuclei were segmented using DAPI signal. For HeLa cells, γH2A.X integrated signal intensity, 53BP1 foci, and PARP1 foci were detected and related to parent nuclei. For HCT116 cells, γH2A.X and 53BP1 foci were detected and related to parent nuclei. Quantified intensity and foci values were plotted in GraphPad Prism 10.0.

Statistical analyses were performed using R software. Quantile Generalized Additive Modeling (qGAM) (37) was employed to model the time-dependent accumulation of DNA damage markers across all cell lines used in the experiments. Baseline differences between the various cell lines were assessed using parametric coefficients derived from the qGAM models, and the statistical significance of these coefficients was evaluated using Wald’s test. To determine the statistical significance of differences in foci accumulation patterns over time among the groups, Likelihood Ratio Tests (LRTs) were conducted. The LRTs compared the full qGAM model, which allows for different time-dependent patterns between the groups, to a null model that assumes all groups follow the same pattern. A Bonferroni test for multiple comparisons was applied when comparing the HCT116 complemented cell lines. A p-value less than 0.05 was considered indicative of statistical significance.

### Colony formation assays

For colony formation assays under DNA damage, Control and ΔPHRF1 (Figure 6) or Control+EV, ΔPHRF1+EV, ΔPHRF1+3XFlag-PHRF1^WT^, and ΔPHRF1+3XFlag-PHRF1^P221L^ (Figure 7) HCT116 cells were harvested at approximately 70% confluence, counted, and seeded into 6-well plates at a density of 500 cells per well in 2 mL in biological triplicate. Cells were allowed to adhere at 37°C overnight. Then old media was replaced with fresh media containing 0, 0.375, 0.75, and 1.5 µg/mL zeocin. Colonies were allowed to for undisturbed over 9 days. Cells were then washed in 1XPBS, fixed in 4% formaldehyde, and stained with 0.5% crystal violet. After washing off residual crystal violet and drying, each well was imaged and growth area was calculated using ImageJ. Then growth data was at each concentration of zeocin was normalized to the growth observed in untreated wells within each cell line. This data was plotted in GraphPad Prism 10, using a Kruskal-Wallis test to determine statistical significance.

## RESULTS

### The PHD finger of PHRF1 binds the N-terminus of histone H3

Our previous examination of the PHD finger domain-containing family for their histone interactions identified the PHD finger of PHRF1 as an H3K4me0 reader (11). This result was unexpected, as a previous study reported this PHD finger to associate with H3K36me3 (18). To address this conflict we started from an *in silico* structural homology model, creating a structural homology model of the PHD finger of PHRF1 bound to the first eight residues of histone H3 using Alphafold-Multimer (Figure 1A and 1B) (19). As shown in Figure 1B, the PHRF1 PHD finger formed a pocket that was found to primarily engage the extreme N-terminus of H3 (residue A1) as well as the side chain of H3K4, which fits into a long, aliphatic groove. This finding is consistent with how other PHD fingers associate with H3K4me0 (38).

Further examination of the aforementioned structural homology model showed that the sidechain of H3A1 fits directly into a pocket formed by residue P221 (Figure 1C). P221 is notable, as this proline is reported in the COSMIC database to be mutated to leucine in multiple cancers (39). To determine if P221L would disrupt PHRF1:H3 interaction, we generated wild-type or P221L mutated forms of the PHRF1 RING-PHD region as GST-tagged recombinant fusions (GST-PHRF1^RP^ and GST-PHRF1^RP(P221L)^, respectively) for a range of *in vitro* biochemical experiments. Additionally, we also generated a version that possessed just the PHD finger of PHRF1 (GST-PHRF1^PHD^) (Figure 1D). These proteins were applied on our histone PTM peptide microarrays, which contain over 300 differentially-modified histone peptides. These analyses confirmed our previous finding that the PHD region of PHRF1 binds to H3 peptides that are unmodified at H3K4 (Figure 1E top and middle images); quantification of binding is found in Figure 1F and Supplementary File 1. Importantly, the GST-PHRF1^RP(P221L)^ fusion showed virtually no binding to any histone peptide on the microarray (Figure 1E bottom, 1F, and Supplementary File 1), validating the importance of P221 in PHRF1 for H3 interaction. We further characterized this H3 interaction quantitatively by probing the binding of our PHRF1 constructs with a C-terminally 5-FAM-labeled peptide comprised of residues 1-20 of histone H3 (H3 1-20 (FAM)) using fluorescence polarization. Again, both GST-PHRF1^RP^ and GST-PHRF1^PHD^ bound to H3 1-20(FAM) to similar degrees (KD = 3.77 ± 0.66 µM, = 2.60 ± 0.29 µM, respectively), while there was little to no observable binding with GST-PHRF1^RP(P221L)^ (Supplementary Figure S1A).

As histone peptides do not always recapitulate how readers engage the physiological substrate of chromatin – i.e., the nucleosome – we next sought to determine the histone binding preferences of PHRF1 in the context of nucleosomes (40–42). Accordingly, we analyzed our GST-PHRF1 fusion proteins using dCypher, a proximity-binding assay optimized for nucleosome studies (Supplementary Figure S1B). Examination of the wild-type PHD-containing PHRF1 fragment (GST-PHRF1^RP^) confirmed PHD finger-H3 tail binding and further, that this PHD fingers does not interact with other histone tails or any methylated lysines. However, it is notable that adjacent acylation on H3 (e.g., lysine 9ac/cr) increased the interaction of PHRF1 with H3 (Supplementary Figure S1B). This observation is fully consistent with studies by our lab and others that show H3 acylation reduces histone tail-DNA interaction thereby increasing the dynamics and accessibility of the H3 tail for the read-write machinery (22, 43). We also observed heightened binding between GST-PHRF1^RP^ and H3K9me3 nucleosomes versus unmodified nucleosomes as well—an unexpected interaction for a PHD finger. We further validated these dCypher findings using solution-based nucleosome pulldown assays, which again confirmed the PHD finger of PHRF1 in both GST-PHRF1^RP^ and GST-PHRF1^PHD^ constructs. These proteins also exhibited greatly reduced binding to H3K4 methylated nucleosomes relative to unmodified nucleosomes. While the alphascreen assay showed increased binding with the PHD finger of PHRF1 and H3K9me3 nucleosomes, we did not observe any difference in binding between unmodified and H3K9me3 nucleosomes with the pulldown assay. This discrepancy may be an effect of the difference in the nature of the experiment (proximity-based versus traditional liquid immobilized binding assays). Finally, our studies also confirmed that PHRF1-nucleosomal binding was dependent on the intact PHD finger, as the P221L mutation abolished all nucleosome interaction. We also note that while an earlier study reported PHRF1’s PHD to read and bind to H3K36me3 (18), our alphascreen and pulldown assays (Supplementary Figure S1B and Figure 1G) show no difference in PHRF1 binding to H3 unmodified versus H3K36me3 nucleosomes, indicating that any observed H3K36me3 binding in the earlier study was likely due to sustained interaction at the H3 N-terminus. Finally, like with peptides, we quantified PHRF1 binding to unmodified H3 on nucleosomes through biolayer interferometry (BLI). GST-PHRF1^RP^ exhibited similar binding affinity towards its binding partner on nucleosomes as on peptides (KD = 1.01 ± 0.014 µM, Supplementary Figure S1C; similar to that measured by fluorescence polarization with H3 1-20 (FAM) peptide)). Again, GST-PHRF1^RP(P221L)^ showed no detectable binding to nucleosomes (Supplementary Figure S1D).

By utilizing orthogonal binding approaches, structural homology modeling, and modified histone peptide and nucleosome libraries, our results show PHRF1’s PHD to have distinct preference for binding the extreme N-terminus of H3 and that this binding is crucially controlled by one residue that is also reported to be mutated in cancer, PHRF1 P221.

### BioID reveals PHRF1 interactions with DNA- and RNA-associated proteins

Having analyzed how the reader domain of PHRF1 binds to histones *in vitro*, we next sought to determine what kind of interactions the full-length PHRF1 protein makes in cells. This was necessary in order to determine the functional role the histone reader domain has in PHRF1. To accomplish this, we conducted a BioID mass spectrometry assay in HeLa cells by labeling exogenous PHRF1 with the miniTurbo biotin ligase enzyme. A miniTurbo-PHRF1 fusion was expressed in HeLa cells and upon addition of biotin, associated proteins were pulled down with streptavidin and identified via mass spectrometry (Figure 2A). NLS-miniTurbo- and miniTurbo-tagged GFP were used as negative controls. This interactome revealed around 150 proteins significantly enriched with PHRF1 versus the GFP negative controls – these largely being proteins involved in splicing, transcription, DNA damage response, and cell cycle regulation (Figure 2B; Supplementary File 2). We also note that as a positive control, PHRF1 itself was identified as one of the most highly enriched proteins in this assay (red circle, Figure 2B).

**Figure 2.**
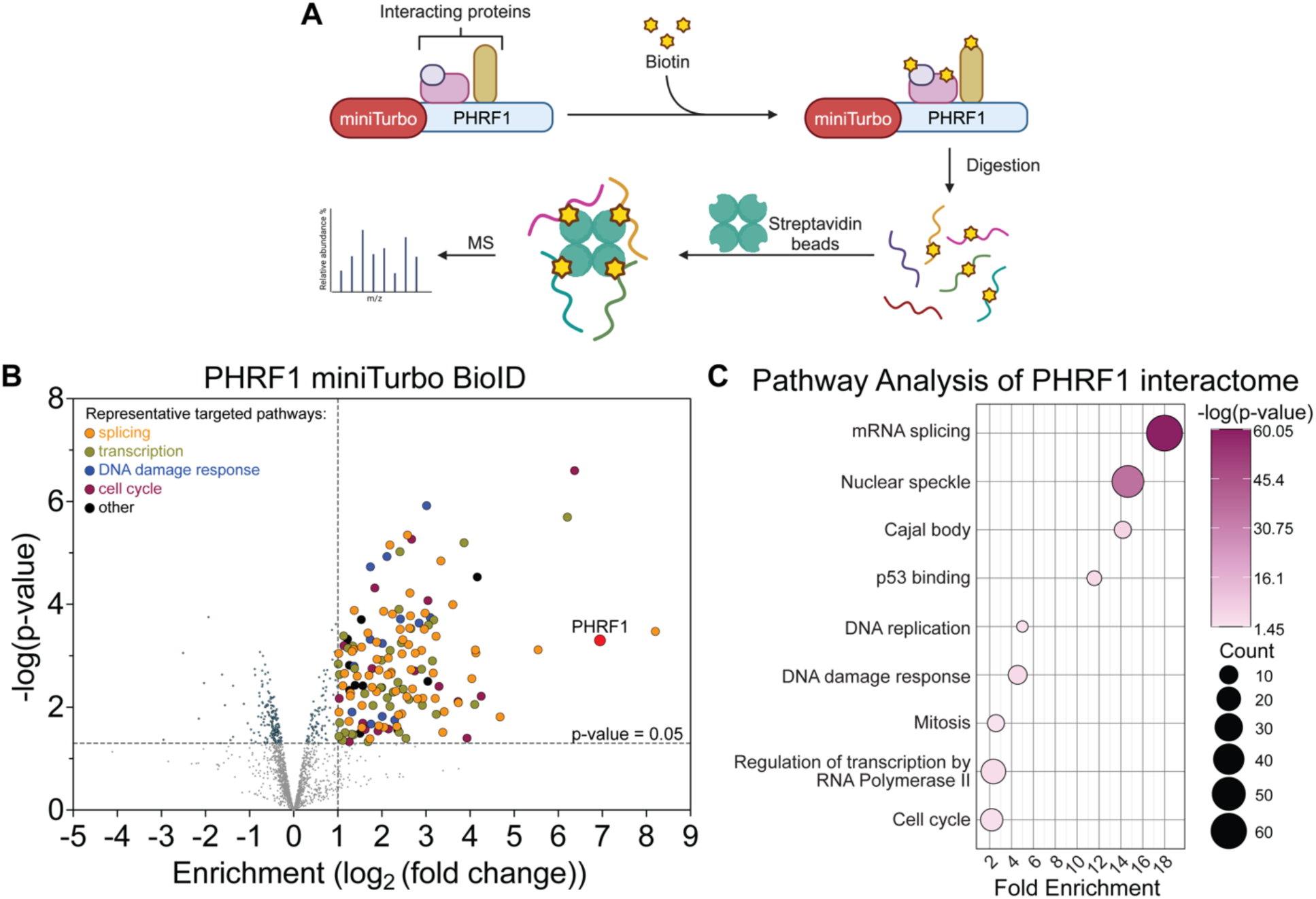
PHRF1 associates with splicing and DNA damage response (DDR)-related proteins *in vivo*. **A.** Schematic of miniTurbo-PHRF1 and the experimental approach for the proximity-based biotinylation assay. **B.** Volcano plot of the miniTurbo-PHRF1 BioID MS analysis, with splicing-relayed proteins shown in orange, transcription-related proteins shown in mustard green, DDR-related proteins shown in blue, cell cycle-related proteins shown in purple, PHRF1 itself shown in red and labeled, and other significant hits shown in black. The x-axis shows enrichment (log2(fold-change)) of proteins in miniTurbo-PHRF1 samples compared to miniTurbo-GFP and NLS-miniTurbo-GFP negative controls; the y-axis shows significance of enrichment (-log(p-value)). Dashed lines at log2(fold-change) = 1 and p-value = 0.05 signify cutoffs for statistical significance. Experiment done in biological duplicate. **C.** Pathway analysis of significant hits from the BioID assay using DAVID. The x-axis is fold enrichment, GO terms are ordered in decreasing fold enrichment along the y-axis. The size of each bubble represents gene count and statistical significance is shown as a purple to pink gradient with all terms having a p-value < 0.05.

Further analysis of the gene ontology terms for enriched targets shed light on additional and likely mechanisms through which PHRF1 may function in the nucleus. For example, PHRF1 was associated with proteins linked to nuclear speckles and Cajal bodies, hubs for splicing and transcriptional activity, and p53 binding, a hallmark for cellular DDR signaling (44, 45) (Figure 2C; Supplementary File 3). These findings are consistent with the predicted biology of PHRF1, as it contains an SRI domain that would make this protein co-transcriptional and associated with splicing machinery (46, 47). DNA damage-associated proteins is also consistent with previous studies that link PHRF1 to DNA repair (18). Consistent with the gene ontology, we confirmed the association of PHRF1 with proteins representative of these processes through co-immunoprecipitation, which showed that PHRF1 associates with transcription (RNA Pol II), splicing (SRSF1), DDR (PARP1), and chromatin itself (H3) (Supplementary Figure S2). Taken together, our comprehensive proteomic analysis of PHRF1’s interactome highlights a number of important pathways and binding partners that provide a bird’s eye view of PHRF1’s regulatory roles in the cell.

### PHRF1 regulates the transcription of DDR- and splicing-associated genes

Given PHRF1 interacts with a large number of chromatin- and transcription-associated proteins, we next asked what genes would be regulated by PHRF1. By using an sgRNA targeting endogenous PHRF1 and CRISPR technology, we generated a PHRF1 knockout in HeLa cells (Figure 3A inset; see Materials and Methods for details) and conducted a deep read (80 million reads/sample) RNA-seq experiment, comparing steady state transcription of genes in ΔPHRF1 cells versus control cells. After applying an FDR cutoff of 0.0001, 2,178 genes were upregulated compared to 2,598 genes being downregulated in ΔPHRF1 cells as compared to control (Figure 3A; Supplementary File 4). Ingenuity Pathway Analysis (IPA) (48) was used to amalgamate the patterns of significantly differentially expressed genes and the extent of their up or downregulation to understand which cellular pathways are regulated by PHRF1. Interestingly, many of the most upregulated pathways involve cell cycle checkpoint regulation and DDR while significantly downregulated pathways involve transcription and splicing (Figure 3B; Supplementary File 5 for full list). Considering the BioID results, these findings imply that PHRF1 may be affecting splicing and DDR through direct protein-protein interaction as well as through the regulation of genes in these pathways.

**Figure 3.**
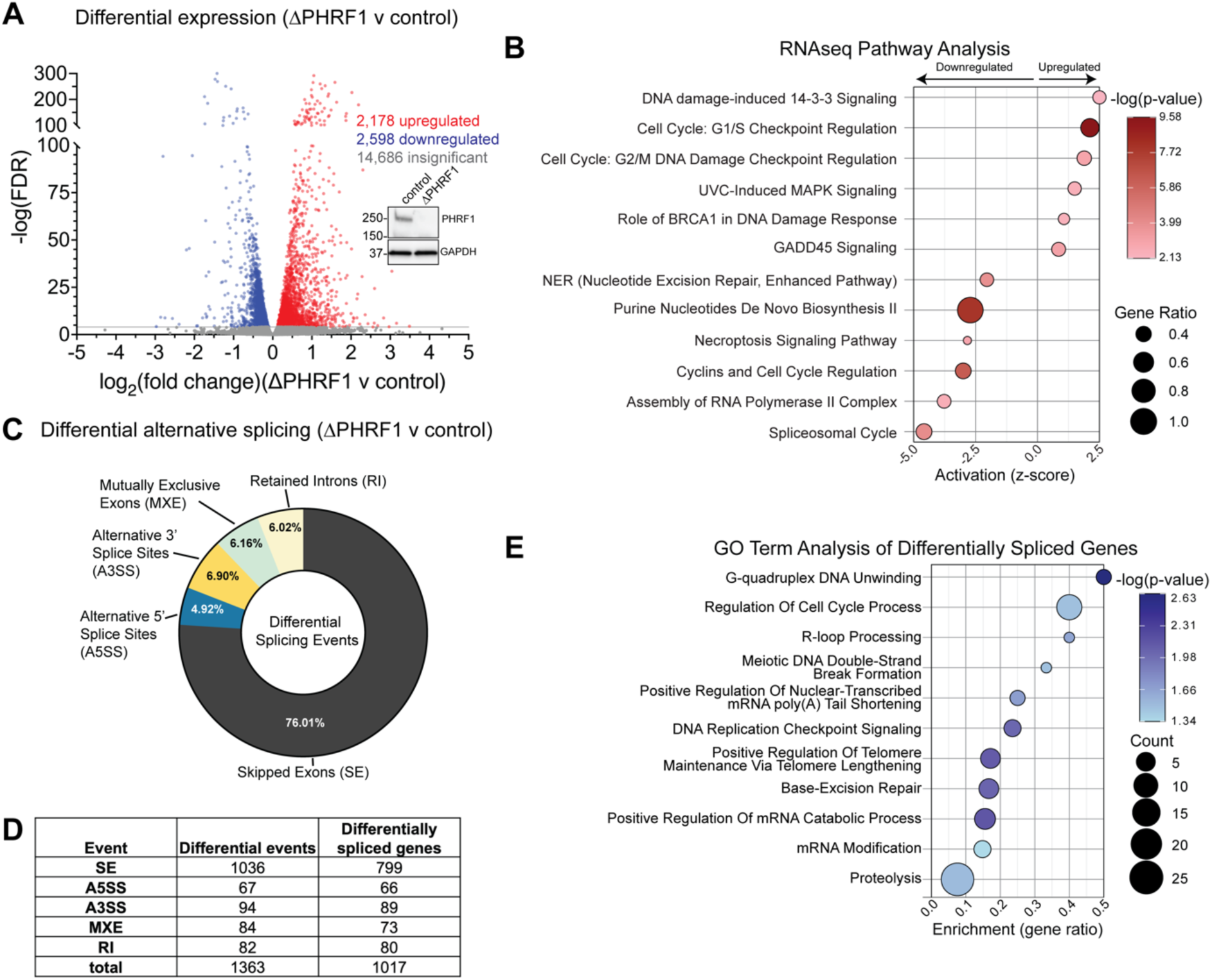
Loss of PHRF1 affects the expression of DDR and splicing-related pathways in HeLa cells. **A.** Volcano plot comparing the expression of genes in ΔPHRF1 and control HeLa cells. After applying an FDR cutoff of 0.0001, significant hits that are upregulated in ΔPHRF1 cells are shown in red and significant hits that are downregulated in ΔPHRF1 cells are shown in blue. The x-axis shows fold change in ΔPHRF1 cells over control cells ((log2(fold-change)) and the y-axis shows statistical significance (-log(FDR)). Inset: western blot analysis of PHRF1 protein levels in control and ΔPHRF1 HeLa cells. Experiments were done in biological triplicate. **B.** Pathway analysis of statistically significant changes in gene expression (FDR <0.0001) using Ingenuity Pathway Analysis (IPA). The x-axis shows activation scores (z-score) of canonical pathways listed on the y-axis, with a positive z-score meaning upregulation and a negative z-score meaning downregulation. Bubble size represents gene ratio and each pathway’s statistical significance is indicated by a color gradient from brown to white. **C.** Summary pie chart of total alternative splicing identified by rMATS from the significant reads from panel **A** after filtering out events with an FDR of > 0.05 and Δ4′ values of <0.01. **D.** Summary table of events represented in panel **C**. Specifically, the number of differential alternative splicing events as well as the number of corresponding genes are indicated for each of the five annotated splicing event types. **E.** GO term analysis of differentially alternatively spliced genes from panel **D** using Enrichr and the GO Biological Process 2023 database. The x-axis shows enrichment of each pathway as gene ratio, the bubble sizes represent gene count, and significance is denoted by a color gradient from blue to white.

Particularly, a large portion of hits identified in the BioID experiments were splicing factors (Figure 2B and 2C; Supplementary Files 2 and 3) and pathway analysis of the RNA-seq data reveals the splicing pathways to be severely downregulated (Figure 3B; Supplementary File 5). Thus, we leveraged the high depth with which the RNA-seq was performed to quantify transcripts with high confidence and identify differential alternative splicing events in ΔPHRF1 cells through replicate multivariate analysis of transcript splicing (rMATS) (29). rMATS enabled us to detect and quantify five types of alternative splicing events—skipped exons (SE), alternative 5’ and 3’ splice sites (A5SS and A3SS, respectively), mutually exclusive exons (MXE), and retained introns (RI)— by analyzing both exon junctions and exon bodies. As shown in Figure 3C and Supplementary File 6, the majority of differential alternative splicing events observed upon PHRF1 knockout were SEs (76.01%), followed by A3SS (6.90%), MXE (6.12%), RI (6.02%), and A5SS (4.92%) (FDR < 0.05, ΔΨ (inclusion difference) > 0.1). This represented 1,363 differential alternative splicing events which comprised of 1,017 unique genes (Figure 3D). To understand whether there was

any significance to the genes whose alternative splicing was affected by the loss of PHRF1, we performed GO-term analysis on the 1,017 genes identified from rMATS using Enrichr (31) and the GO Biological Process 2023 database. Once again, processes related to DDR, particularly involving DNA-RNA interactions such as R-loop processing, were enriched (Figure 3E; Supplementary File 7). Our rMATS analysis also identified a statistically significant increase in the inclusion levels of A5SS and RI events in ΔPHRF1 cells compared to control cells (p-value < 0.05; Figure S3).

### ΔPHRF1 HeLa cells exhibit defective molecular DNA damage response

A recurring theme in our proteomic and genomic analyses of PHRF1 is the enrichment of DDR pathways – a finding consistent with work by others who have linked PHRF1 to DDR (18). However, the understanding of the precise roles of PHRF1 in DDR remains elusive. Accordingly, we developed a DNA damage recovery assay in which control and ΔPHRF1 cells were treated with 50 µg/mL zeocin, a double-strand break (DSB)-inducing drug (49–51), for 30 minutes. The drug was then washed out and cells were collected along the washout/recovery phase for immunoblotting or probed for markers of DNA damage via immunofluorescence (IF) (Figure 4A). Interestingly, PHRF1 protein levels in control cells increase during the recovery phase compared to the untreated/undamaged cells (Supplementary Figure S4A). γH2A.X (H2A.X S139ph) was used as a marker for DNA damage in the IF experiments, and as can be seen through representative images, during the recovery phase of the experiment, ΔPHRF1 cells show a much higher and sustained accumulation of γH2A.X relative to control cells (Figure 4B). This phenomenon was quantified by measuring γH2A.X intensity/nucleus in Cell Profiler (Figure 4C) and them performed quantile general additive modeling (qGAM) to determine their statistical significance (Figure 4D; Supplementary Table S1). The baseline γH2A.X intensity/nucleus for much lower in control cells versus ΔPHRF1 as exhibited by the Parametric Coefficient value of - 345.25 (Supplementary Table S1). We then used a Likelihood Ratio test (LRT) comparing the full qGAM model, which allows for different time-dependent patterns between two groups/cell lines, with the null model, which assumes both groups follow the same pattern. The LRT indicated that the full model was a much better fit for the data than the null model (Supplementary Table S1, LRT p-value 3.26 x10^-10^ This suggests that not only did control cells have much lower levels of γH2A.X during the recovery phase than ΔPHRF1, but γH2A.X accumulation also did not change as dramatically over time in control cells as in ΔPHRF1 cells. We also probed these cells for other hallmarks of DNA damage, namely 53BP1 and PARP1 foci formation (52). Immunofluorescence revealed that 53BP1 and PARP1 foci to form in higher quantities 4 hours post zeocin washout in ΔPHRF1 cells versus control cells (Supplementary Figure S4B-D), consistent with the overall effects observed with γH2A.X accumulation. Taken together, these results clearly indicate that HeLa cells lacking PHRF1 accumulate more DNA damage than cells expressing PHRF1, and that the kinetics of repair may also be deficient in ΔPHRF1 cells.

**Figure 4.**
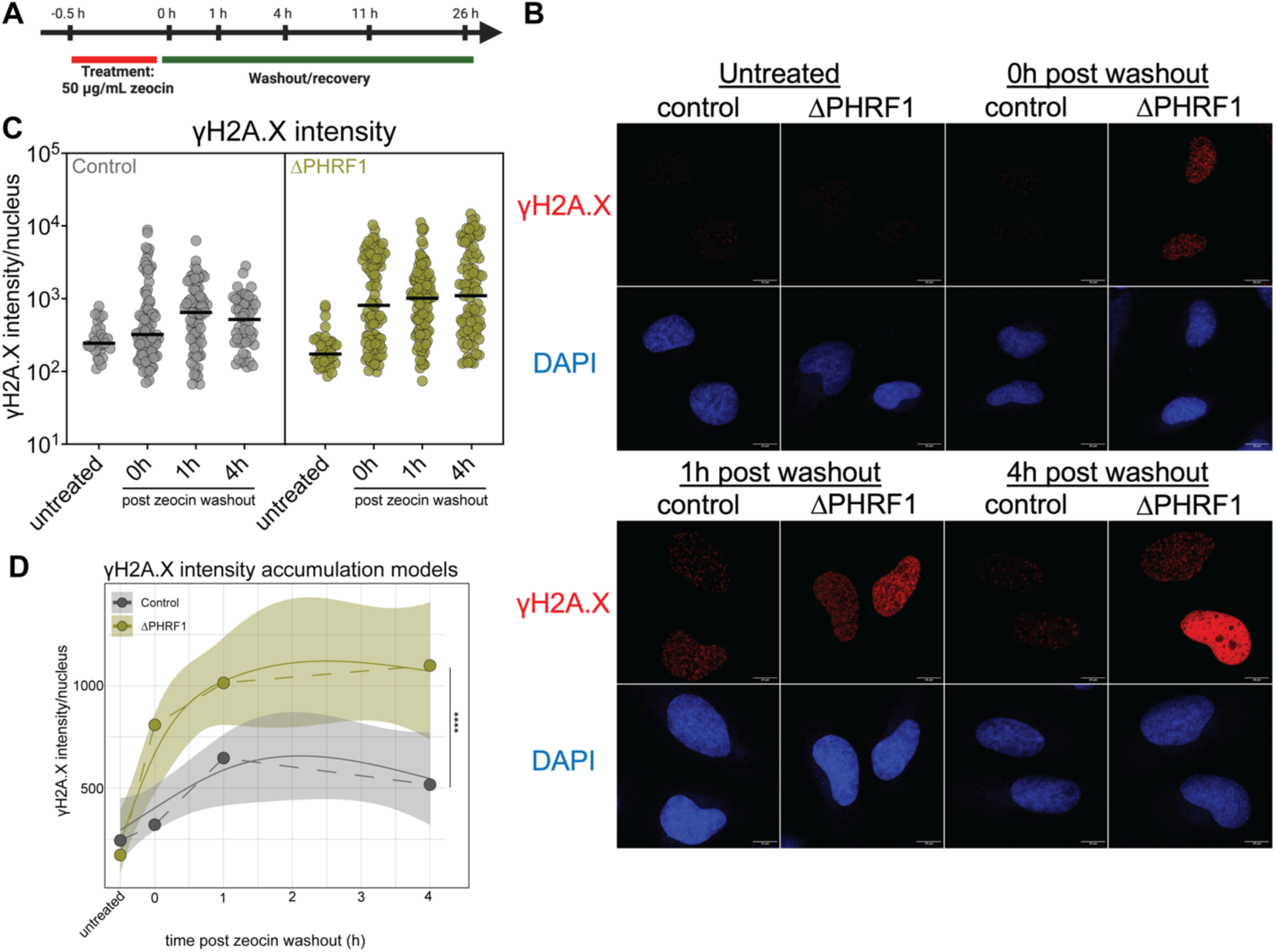
Loss of PHRF1 results in a defective DNA damage response. **A.** Experimental schematic of zeocin DDR assay in HeLa cells. Time points indicate when cells were collected for analysis along the experimental trajectory. **B.** Representative confocal immunofluorescence of control and ΔPHRF1 cells stained with anti-γH2A.X (red) antibody and DAPI (blue) at indicated time points along the zeocin DDR assay. Scale bar is 10 μm. **C.** Quantitative analysis of γH2A.X intensity/nucleus in control (gray) and ΔPHRF1 (mustard green) cells in the zeocin DDR assay. Median values are shown with solid black lines. **D.** Quantile general additive modeling (qGAM) of γH2A.X signal accumulation over time from panel **C** for control (gray) and ΔPHRF1 (mustard green) cells. Median values are represented as solid, colored circles connected by dashed lines. The qGAM generated models are represented with smooth lines and shaded in 95% confidence intervals. Statistical significance was determined using a Wald test. **** = p-value < 0.0001. See Supplementary Table S1 for full statistical analysis.

### PHRF1 is highly expressed in colorectal cancer cells and exhibits similar transcriptional effects in HCT116 as seen in HeLa cells

Improper or deficient cellular DNA damage response is a hallmark of cancer cells and often exploited for therapeutic advantage in the treatment of cancer (44, 53, 54). As such, we wanted to determine whether the heightened DNA damage phenotype we observed in ΔPHRF1 HeLa cells would also be true in other, more cancer-relevant model systems (29). To begin to address this, we compared PHRF1 expression levels across different cancer types from the Cancer Cell Line Encyclopedia (CCLE) (Figure 5A). From this analysis, PHRF1 was found to be most highly expressed in cancers derived from blood, digestive, and reproductive tissue types. Consistent with this, a western blot analysis of a panel of representative model cancer cell lines showed PHRF1 protein levels to be highest in cell lines from digestive tissues, most notably HCT116 cells (Figure 5B). Additionally, high expression of PHRF1 is also linked to poorer survival across a wide range of patient cancers (Supplementary Figure S5).

**Figure 5.**
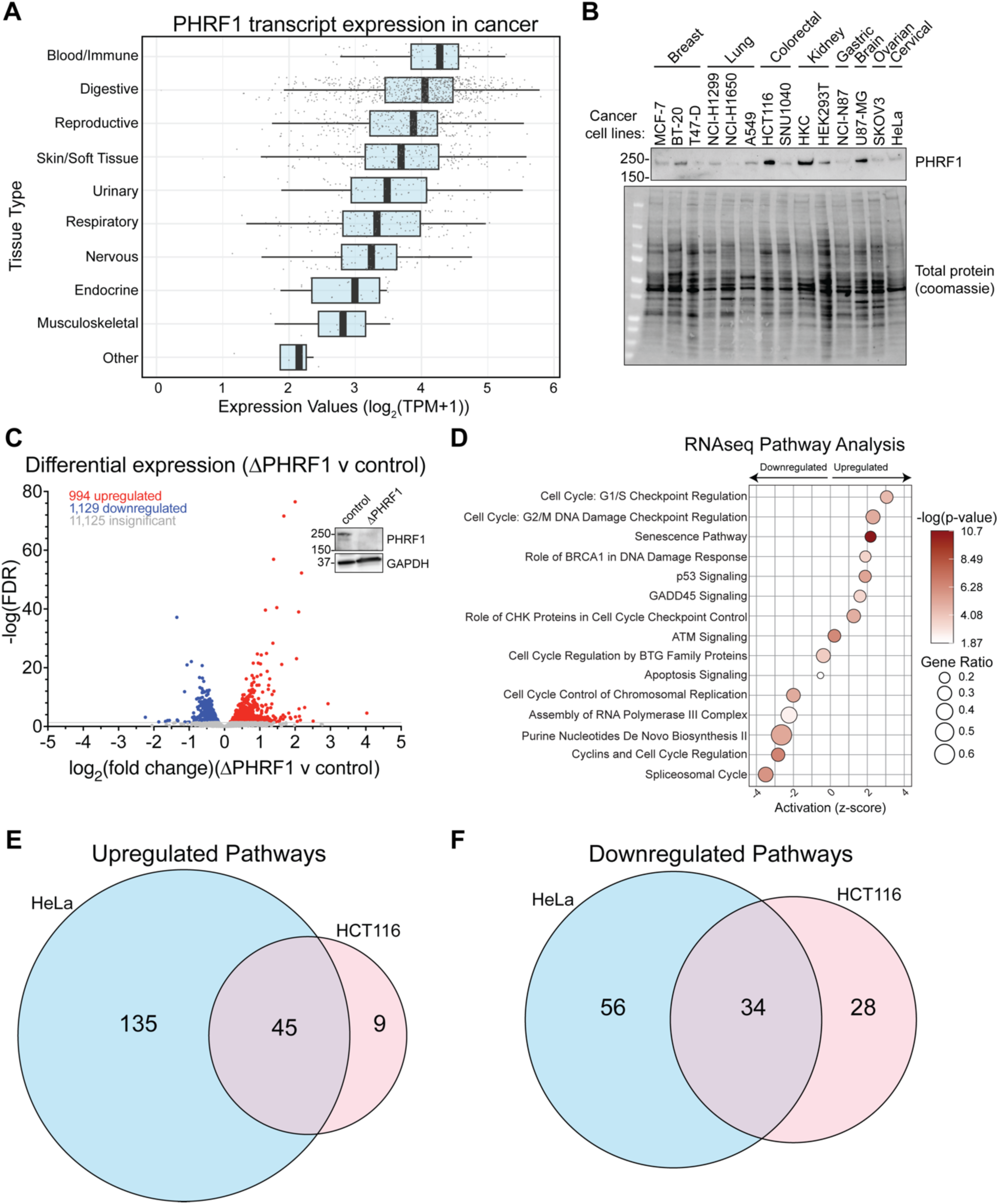
PHRF1 is highly expressed in HCT116 colorectal cancer cells and is required for appropriate expression of genes involved in splicing and DDR-related pathways. **A.** Boxplots illustrate distribution of PHRF1 expression levels (log_2_(TPM+1)) in cancer cells from the Cancer Cell Line Encyclopedia (CCLE) collapsed into the indicated tissue types on the y-axis. The central line in each box represents the median. Each cell line’s individual PHRF1 expression is represented as a gray circle. **B.** Western blot of PHRF1 protein levels in the indicated cancer cell lines and their corresponding cancer type category. **C.** Volcano plot comparing the expression of genes in ΔPHRF1 and control HCT116 cells. After applying an FDR cutoff of 0.05, significant hits that are upregulated in ΔPHRF1 cells are shown in red and significant hits that are downregulated in ΔPHRF1 cells are shown in blue. The x-axis shows fold change in ΔPHRF1 cells over control cells ((log_2_(fold-change)) and the y-axis shows statistical significance (-log(FDR)). Inset: western blot analysis of PHRF1 protein levels in control and ΔPHRF1 HCT116 cells. Experiments were done in biological triplicate. D. Pathway analysis of statistically significant changes in gene expression (FDR <0.05) using IPA, as in Figure 3B. Bubble size represents gene ratio and each pathway’s statistical significance is indicated by a color gradient from brown to white. **E, F.** Venn diagram comparison of the number of upregulated (**E**) and downregulated (**F**) pathways identified from HeLa and HCT116 RNAseq experiments.

Given HCT116 cells mimic the genetic and phenotypic characteristics of colorectal cancer (30) and are well-suited to study genomic instability and DDR in cancer (55), we generated a PHRF1 knockout in HCT116 cells and first conducted an RNA-seq experiment (Figure 5C; Supplementary File 8). After applying an FDR cutoff of 0.05, we observed 994 genes to be upregulated and 1,129 genes to be downregulated in ΔPHRF1 cells compared to control cells. A more stringent cutoff was applied to the HeLa dataset versus HCT116 cells because that experiment was performed with much greater read depth than the HCT116 dataset. Ingenuity Pathway Analysis was performed on this list of differentially expressed genes (DEGs) and, similar to our observations in HeLa cells, pathways related to cell cycle checkpoint regulation and DDR were found to be upregulated, while splicing-related pathways were significantly downregulated (Figure 5D; Supplementary File 9). Indeed, when we compared the up and downregulated pathways identified in the HeLa and HCT116 RNA-seq experiments, we found that a majority of up and downregulated pathways in HCT116 cells were also similarly regulated in HeLa cells (Figure 5E and 5F), implying the role of PHRF1 fundamentally conserved across diverse tissues.

### Loss of PHRF1 results in significant defects in cellular DDR and enhanced sensitivity to DSB-causing agents

Having established that loss of PHRF1 caused similar changes in gene expression in HCT116 cells as in HeLa cells, we asked how ΔPHRF1 HCT116 cells responded to DNA damage stress compared to control cells using IF. In a similar DNA damage recovery assay as the one performed in HeLa cells (Supplementary Figure S6A), ΔPHRF1 and control cells were co-stained with antibodies for γH2A.X and 53BP1 (red and green, respectively, Figure 6A). Unlike HeLa cells, where only pan-nuclear γH2A.X staining was observed, HCT116 showed distinct γH2A.X foci along with 53BP1 foci (yellow foci in “Merge” panels, Figure 6A). This indicates that HeLa cells may undergo additional non-DSB-related stress upon zeocin treatment, resulting in broad accumulation of γH2A.X, while in HCT116 the major inducer of DNA damage is the formation of DSBs by zeocin, resulting in discrete γH2A.X foci. This also supports the use of HCT116 cells to study the role of PHRF1 in DDR further. Interestingly, while in control HeLa cells PHRF1 levels increased during the washout/recovery phase, they appeared to be relatively stable in HCT116 cells (Supplementary Figure S6B). As before, once γH2A.X and 53BP1 foci accumulation were quantified (Figure 6B and 6C), we performed qGAM analysis. This analysis showed that there were significant baseline differences between ΔPHRF1 and control for both γH2A.X foci (Figure 6D) and 53BP1 foci (Figure 6E) overall. Specifically, parametric coefficients for both γH2A.X and 53BP1 foci were significantly lower in control cells versus ΔPHRF1 cells, -1.9417 and -3.4801, respectively (Supplementary Table S2), consistent with results observed in HeLa cells. Using the Likelihood Ratio Test (LRT), we also observed a significant difference in the foci accumulation patterns of γH2A.X and 53BP1 over the recovery phase of this assay (Supplementary Table S2, LRT p-values). Both observations show that ΔPHRF1 cells exhibit heightened and differential response to DNA damage than control HCT116 cells, in agreement with results from HeLa cells. We further tested whether these changes in DDR on the molecular level had implications for cell viability. Colony formation assays under a titration of zeocin concentration showed that zeocin was cytotoxic to a much higher degree for ΔPHRF1 cells versus control HCT116 cells (Figure 6F, 6G, and Supplementary Figure S6C).

**Figure 6.**
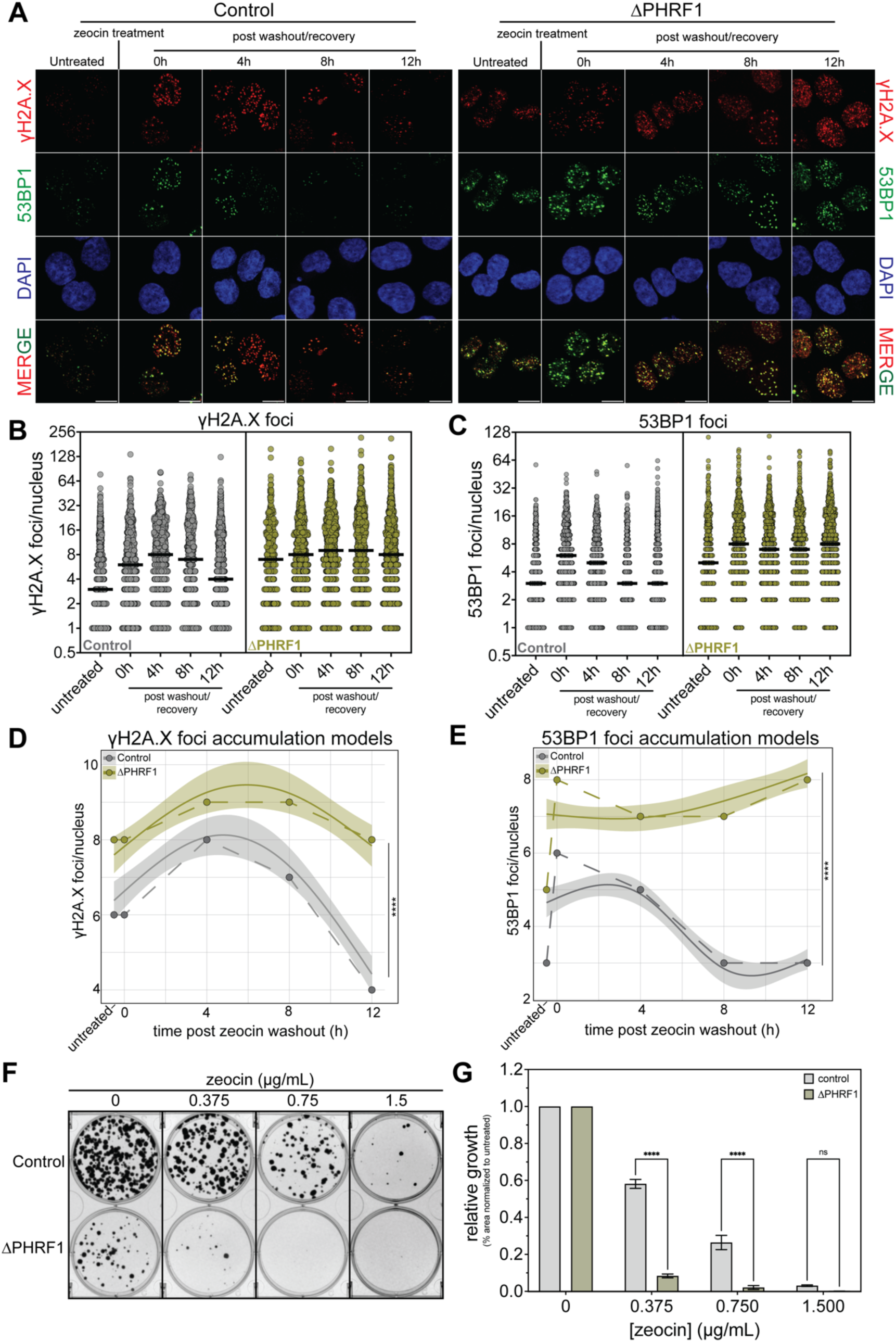
Loss of PHRF1 results in defective cellular DDR and cell survival in HCT116 cells. **A.** Representative confocal immunofluorescence of control and ΔPHRF1 cells stained with anti-γH2A.X (red), anti-53BP1 (green) antibodies, and DAPI (blue) at indicated time points along the zeocin DDR assay, as described in **Supplementary Figure S6A**. The scale bar is 10 μm. **B, C.** Quantitative analysis of γH2A.X (**B**) and 53BP1 (**C**) foci/ nucleus in control (gray) and ΔPHRF1 (mustard green) HCT116 cells. Median values are shown with solid black lines. **D, E.** qGAM of γH2A.X (**D**) and 53BP1 (**E**) foci/nucleus accumulation over time for control (gray) and ΔPHRF1 (mustard green) cells. Median values are represented as solid, colored circles connected by dashed lines. The qGAM-generated models are represented with smooth lines and shaded in 95% confidence intervals. Statistical significance was determined using a Wald test. **** = p-value < 0.0001. See **Supplementary Table S2** for full statistical analysis. **F.** Representative images of a colony formation assay (CFA) comparing growth in control and ΔPHRF1 HCT116 cells at the indicated concentrations of zeocin. This experiment was conducted in biological triplicate. **G.** Quantification of growth from the CFA in panel **F**, comparing control (gray) and ΔPHRF1 (mustard green) HCT116 cells. Statistical significance was determined through a Kruskal-Wallis test; **** = p-value <0.0001, ns = not significant.

### PHRF1-histone reading is important for proper cellular DDR

We next sought to determine if the histone reading function of PHRF1 was involved in the DDR phenotype we found in ΔPHRF1 HCT116 cells. Accordingly, we performed a complementation experiment and expressed either a wild-type (3XFlag-PHRF1^WT^) or PHD mutant form (3XFlag-PHRF1^P221L^) of PHRF1 into ΔPHRF1 HCT116 cells (Supplementary Figure S7A). We confirmed that expression of these constructs was similar to that of endogenous PHRF1 in control HCT116 cells (Supplementary Figure S7B) and cell growth was monitored for four HCT116 complimented cell lines: Control+Empty Vector (EV), ΔPHRF1+EV, ΔPHRF1+3XFlag-PHRF1^WT^, and ΔPHRF1+3XFlag-PHRF1^P221L^. Under normal growth conditions, we observed that ΔPHRF1+EV cells (mustard green) showed the slowest growth relative to Control+EV (gray) (Supplementary Figure S7C). This was in contrast to what was observed with the expression of the WT PHRF1 construct (green), which largely rescued the growth defects observed in ΔPHRF1 cells. Our analysis of the P221L-containing PHRF1 construct showed this form of PHRF1 was also capable of partially rescuing the growth phenotype found in ΔPHRF1 cells, and further, showed slightly better growth in the early phase of the experiment relative to exogenously expressed WT (red versus green). The better growth of the P221L-containing cells may be explained by the slightly elevated expression levels of the P221L construct compared to the WT construct (Supplementary Figure S7B). Therefore, under standard growth conditions, these data reveal that the PHD finger of PHRF1 is dispensable for normal cellular growth. These cellular growth findings are in stark contrast to our findings (described below) that reveal a role for the PHD finger of PHRF1 in DDR.

We next sough to determine what role, if any, the PHRF1 PHD finger might have in DDR. Similar to Figure 6F and 6G, we performed colony formation assays with our complementation HCT116 cell lines and performed a DNA damage experiment as before (Figure 7A; Supplementary Figure S7D). Quantification of cellular growth revealed that while ΔPHRF1+EV cells grew much slower than Control+EV cells at all concentrations of zeocin (Figure 7B mustard green), this defect was rescued by the expression of 3XFlag-PHRF1^WT^ (green). While we observed a partial rescue of growth at 0.375 µg/mL by both the WT and P221L constructs, it should be noted that such effects were greater with ΔPHRF1+3XFlag-PHRF1^WT^ cells relative to ΔPHRF1+3XFlag-PHRF1^P221L^. Moreover, at higher zeocin concentrations, particularly 0.75 µg/mL, expression of 3XFlag-PHRF1^P221L^ was unable to rescue ΔPHRF1’s growth defect, while the cells expressing exogenous WT PHRF1 were able to do so (mustard green versus red; Figure 7A). This indicates a role for PHRF1 histone reading in DNA damage response.

**Figure 7.**
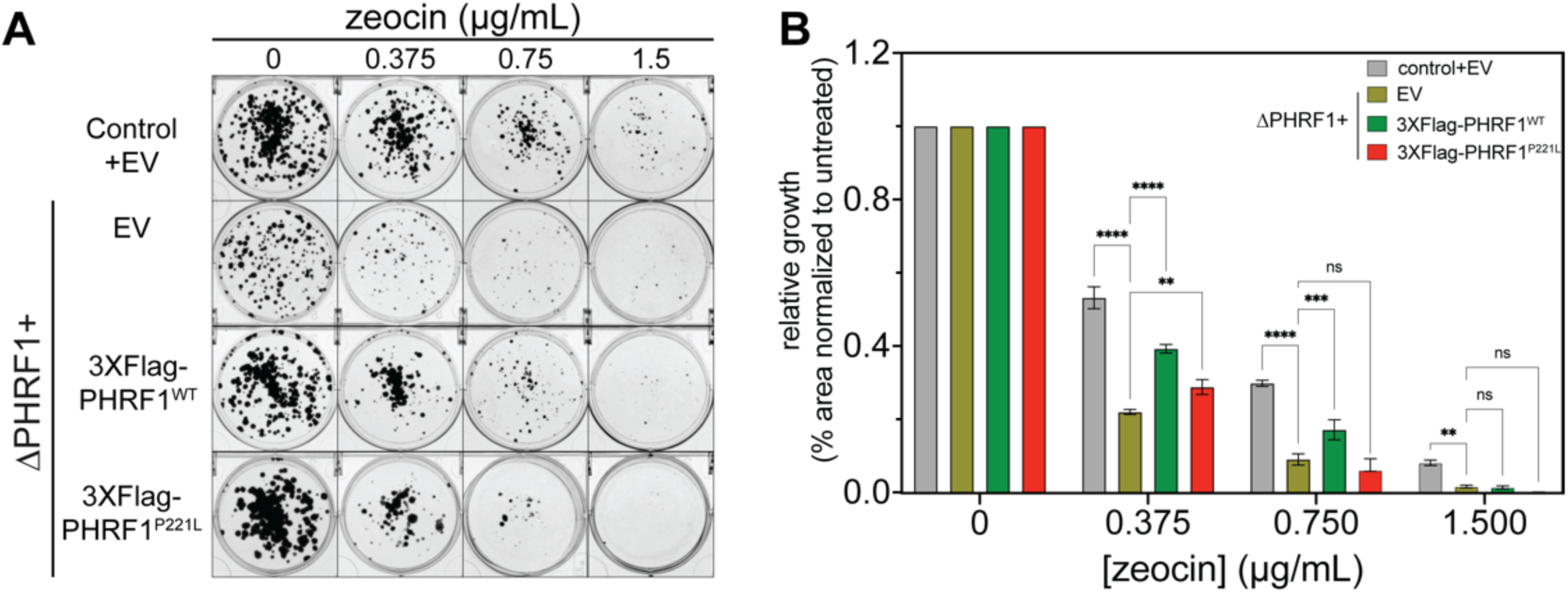
PHD finger binding activity of PHRF1 is required for proper cellular DNA damage response. **A.** Representative images of a colony formation assay (CFA) comparing growth in Control+EV (empty vector) and ΔPHRF1 + indicated complemented HCT116 cells at the indicated concentrations of zeocin. This experiment was conducted in biological triplicate. **B.** Quantification of growth from the CFA in panel **A**, comparing ΔPHRF1+EV (mustard green) cells to Control+EV (gray), ΔPHRF1+3XFlag-PHRF1^WT^ (green), and ΔPHRF1+3XFlag-PHRF1^P221L^ (red) HCT116 cells. Error bars are S.D. Statistical significance was determined through a Kruskal-Wallis test; **** = p-value <0.0001, *** = p-value <0.001, ** = p-value <0.01, * = p-value <0.05, and ns = not significant.

Finally, we performed the same zeocin DNA damage recover assay described in Supplementary Figure S6A with our complemented HCT116 cell lines. We confirmed stable expression of PHRF1 in both cells with endogenous PHRF1 and those complemented with exogenous PHRF1 (Supplementary Figure S8A). Confocal microscopy with γH2A.X and 53BP1 staining revealed significant accumulation of DNA damage foci in Control+EV (Figure 8A), ΔPHRF1+EV (Figure 8B), ΔPHRF1+3XFlag-PHRF1^WT^ (Figure 8C), and ΔPHRF1+3XFlag-PHRF1^P221L^ (Figure 8D) cells after damage and during recovery, showing different patterns among the cell lines. After quantifying γH2A.X (Supplementary Figure S8B) and 53BP1 (Supplementary Figure S8C) foci per nucleus, we performed qGAM analysis as in Figure 6 (Figure 8E-J). This analysis showed that ΔPHRF1+EV cells had significantly higher foci per nucleus compared to Control+EV and ΔPHRF1+3XFlag-PHRF1 ^WT^ cells (see Parametric Coefficients in Supplementary Table S3), indicating increased DNA damage due to PHRF1 loss. Notably, ΔPHRF1+3XFlag-PHRF1^P221L^ cells did not show significant improvement over ΔPHRF1+EV cells (Figure 8G and 8J and Supplementary Table S3), indicating that PHRF1 histone reading is necessary for DNA repair. A Likelihood Ratio Test confirmed that while all cell lines exhibit distinct patterns of foci accumulation compared to ΔPHRF1+EV cells, cells expressing WT PHRF1 show the starkest pattern changes (see LRT p-values in Supplementary Table S3), indicative of a functioning DNA damage response network. In total, our results show an important role of PHRF1 in DNA damage repair that depends on its ability to bind to histones.

**Figure 8.**
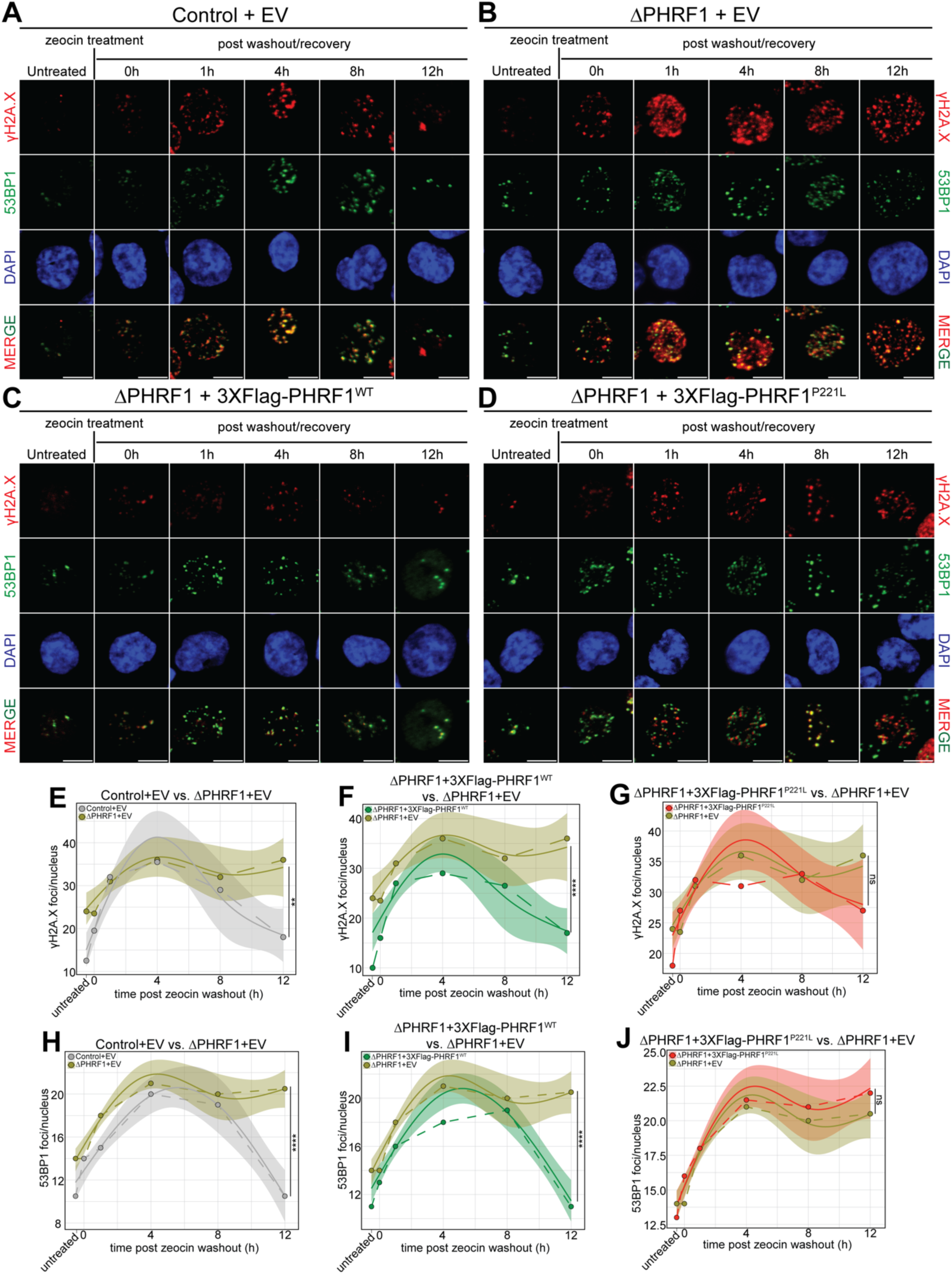
PHD finger binding activity of PHRF1 is required for proper cellular DNA damage response. **A-D.** Representative confocal immunofluorescence of Control+EV (**A**), ΔPHRF1+EV (**B**), ΔPHRF1+3XFlag-PHRF1^WT^ (**C**), and ΔPHRF1+3XFlag-PHRF1^P221L^ (**D**) cells stained with anti-γH2A.X (red), anti-53BP1 (green) antibodies, and DAPI (blue) at indicated time points along the zeocin DDR assay, as described in **Supplementary Figure S6A**. The scale bar is 10 μm. **E-J.** qGAM of γH2A.X (**E-G**) and 53BP1 (**H-J**) foci/nucleus accumulation over time, comparing ΔPHRF1+EV (mustard green) with Control+EV (gray; **E, H**), ΔPHRF1+3XFlag-PHRF1^WT^ (green; **F, I**), and ΔPHRF1+3XFlag-PHRF1^P221L^ (red; **G, J**) cells. Median values are represented as solid, colored circles connected by dashed lines. The qGAM-generated models are represented with smooth lines and shaded in 95% confidence intervals. Statistical significance was determined using a Wald test. **** = p-value < 0.0001, ** = p-value <0.01, and ns = not significant. See **Supplementary Table S3** for full statistical analysis.

## DISCUSSION

Over the last decade, a handful of studies have showcased the biological significance of PHRF1 in cancer biology. These studies showed a role for PHRF1 in tumor growth in breast and lung cancers (12, 13, 15). PHRF1 has also been linked to cancer cell invasion and migration in colorectal and lung cancers (56, 57). Furthermore, one study suggests that PHRF1 may accumulate at sites of DNA damage and is involved in NHEJ-mediated DNA damage repair (18). While these studies collectively underscore PHRF1’s importance as a cancer regulator, they lack consistency and a comprehensive understanding of its biochemical functions, particularly regarding its chromatin interactions and impacts to gene transcription. Given this gap in knowledge, we sought to elucidate the roles of PHRF1 in regulating cancer biology via chromatin-mediated mechanisms. By conducting proteome- and genome-wide experiments alongside rigorous structural and *in vitro* biochemical analyses, we aimed to provide an unbiased and holistic understanding of PHRF1’s functions.

Our biochemical studies demonstrated that the PHD finger of PHRF1 exhibits exquisite specificity for the extreme N-terminus of histone H3, both on peptides and nucleosomes. Importantly, a mutation of a conserved residue within the PHD region, P221 to leucine—a mutation observed in tumors—completely abolished this interaction (Figure 1 and Supplementary Figure S1). Moving into cells, we revealed PHRF1’s interactome, highlighting that this protein predominantly associated with splicing, DNA damage, and cell cycle-associated proteins (Figure 2 and Supplementary Figure S2). This was further corroborated by in-depth genomic and transcriptomic analysis in HeLa and HCT116 cells, again revealing numerous DDR/cell cycle and splicing-related pathways to be significantly perturbed by the loss of PHRF1 expression (Figures 3 and 5 and Supplementary Figure S3). Given the occurrence of DDR pathways in both proteomic and genomic analysis, we asked if PHRF1 influenced how cells responded to DNA damage. Functional assays demonstrated that ΔPHRF1 cells had significantly higher amounts of DNA damage and displayed greater cytotoxicity towards genotoxic agents such as zeocin (phleomycin D1) (50, 51) relative to control cells (Figures 4 and 6). Finally, we showed that complementation of exogenous PHRF1 with a P221L mutation (a proxy for a PHD finger unable to bind H3) in a ΔPHRF1 background could not rescue the DNA damage molecular and growth defects (Figures 7 and 8) although the PHD finger is dispensable for normal cellular growth (Supplementary Figure S7). It is notable that there was a significant difference in the ability of exogenous P221L expressing cells to respond to DNA damage depending on the assays used. In the colony formation DNA damage assay (Figure 7), there is slow and constant accumulation of damage. In this context, the P221L mutation shows only a partial growth defect upon DNA damage. However, this response was observed in a much more exaggerated manner during acute DNA damage (Figure 8), which showed that the P221L mutation completely abrogates the ability of this form of PHRF1 to resolve the DNA damage phenotype (accumulation of damage foci). While both assays reveal an important role for PHRF1’s PHD finger in DDR, we surmise that cells under constant DNA damage find ways to adapt to these conditions and use additional repair mechanisms for survival (however, even in this scenario, the PHD finger and histone reading of PHRF1 is still at least partially required). Nonetheless, these findings highlight the importance of the PHD region and PHRF1’s binding to histones in maintaining proper cellular DDR, a well-established avenue for targeting cancer therapies.

Understanding PHRF1’s involvement in splicing and DDR is crucial for comprehending how it regulates cancer biology. Aberrant splicing is a hallmark of cancer, leading to the production of oncogenic splice variants or the loss of tumor suppressor functions (58, 59). Proteins involved in splicing regulation, when dysregulated, can contribute to tumorigenesis and cancer progression (60, 61). Similarly, defects in DDR pathways result in genomic instability, another hallmark of cancer (54, 62). Our findings suggest that PHRF1 may serve as a pivotal coordinator of these processes through its interactions with RNA Pol II, chromatin, and other associated protein complexes. While we have focused on the impact of PHRF1 deletion or mutation of its chromatin-binding PHD domain, further studies are warranted on the functional contributions and disease relevance of the SRI (Set2-Rpb1 Interacting) and catalytic RING domains.

By conducting such a diverse set of experiments, we were able to confirm, tie together, and expand upon several of the reported finding about PHRF1, such as its involvement in DDR (18), and also shed new light on mechanisms involving processes such as splicing and cell cycle checkpoint regulation. Significantly, we showed that a cancer mutation in the PHD region not only ablates PHRF1’s bind to H3, but also functions as poorly as a ΔPHRF1 background in restoring proper cellular DDR in HCT116 cells, a completely new finding that links PHRF1’s role in chromatin biology to cancer. Based on these findings, we hypothesize that PHRF1 functions in two distinct ways: directly through protein-protein interactions with splicing and DDR machinery and indirectly by altering transcription, impacting pathways important to cancer biology, like DDR, and that PHRF1’s ability to bind H3 is vital to these functions. Having done so, we believe this work has identified new ways in which PHRF1 can be targeted for therapeutic studies in cancer, especially given the effect high expression has on overall cancer survival (Supplementary Figure S5). We have also provided, through comprehensive characterization of PHRF1, new avenues for understanding PHRF1, such as splicing and cell cycle mechanisms. Furthermore, work to study the roles of the RING catalytic domain and the SRI domain in conjunction with the results from this work will provide a better understanding of PHRF1-mediated gene regulation in the future, with obvious implications in cancer research. Our findings emphasize the importance of integrating biochemical, proteomic, and genomic approaches to unravel the complex regulatory networks mediated by chromatin-associated proteins like PHRF1.

## DATA AVAILABILITY

All MS files used in this study were deposited to MassIVE (http://massive.ucsd.edu) and proteomeXchange (http://www.proteomexchange.org/) and were assigned the identifiers MSV000092164 and PXD042986. They can be accessed at ftp://massive.ucsd.edu/MSV000092164/. The password to access the MS files prior to publication is “PHRF1”. The usernames for web access prior to publication is “MSV000092164_reviewer”. The accession numbers for RNAseq datasets can be found on the Gene Expression Omnibus (HeLa: GSE281703, HCT116: GSE281702).

## ACKNOWLEDGEMENTS

We thank members of the Strahl lab members for technical and editorial suggestions. We also thank Dr. Jackson Peterson from Dr. Edward Browne’s lab (UNC School of Medicine) for help with nucleofection experiments; Caroline Fraser from Dr. Ian Davis’ lab (UNC School of Medicine) for initial training on the IncuCyte imaging system; Dr. Amelia McCue from Dr. Brian Kuhlman’s lab (UNC School of Medicine); Dr. Logan Slade from Dr. Jean Cook’s lab (UNC School of Medicine) for advice on microscopy-related experiments; and Dr. Wendy Salmon and the UNC Hooker Imaging Core Facility, which is supported in part by NCI Cancer Center Core Support Grant (2P30CA016086-45) to the UNC Lineberger Comprehensive Cancer Center, for advice and helpful suggestions related to microscopy. Peptides used in this work were synthesized in the UNC Peptide Synthesis Core facility (RRID:SCR_017837).

## REFERENCES

1. Su, Z. and Denu, J.M. (2016) Reading the Combinatorial Histone Language. ACS Chem Biol, 11, 564–574.

2. Strahl, B.D. and Allis, C.D. (2000) The language of covalent histone modifications. Nature, 403, 41–45.

3. Jenuwein, T. and Allis, C.D. (2001) Translating the Histone Code. Science (1979), 293, 1074–1080.

4. Biswas, S. and Rao, C.M. (2018) Epigenetic tools (The Writers, The Readers and The Erasers) and their implications in cancer therapy. Eur J Pharmacol, 837, 8–24.

5. Ali, M., Hom, R.A., Blakeslee, W., Ikenouye, L. and Kutateladze, T.G. (2014) Diverse functions of PHD fingers of the MLL/KMT2 subfamily. Biochimica et Biophysica Acta (BBA) - Molecular Cell Research, 1843, 366–371.

6. Dhar, S.S., Lee, S.-H., Kan, P.-Y., Voigt, P., Ma, L., Shi, X., Reinberg, D. and Lee, M.G. (2012) Trans-tail regulation of MLL4-catalyzed H3K4 methylation by H4R3 symmetric dimethylation is mediated by a tandem PHD of MLL4. Genes Dev, 26, 2749–2762.

7. Migliori, V., Müller, J., Phalke, S., Low, D., Bezzi, M., Mok, W.C., Sahu, S.K., Gunaratne, J., Capasso, P., Bassi, C., et al. (2012) Symmetric dimethylation of H3R2 is a newly identified histone mark that supports euchromatin maintenance. Nat Struct Mol Biol, 19, 136–144.

8. Appikonda, S., Thakkar, K.N. and Barton, M.C. (2016) Regulation of gene expression in human cancers by TRIM24. Drug Discov Today Technol, 19, 57–63.

9. Agricola, E., Randall, R.A., Gaarenstroom, T., Dupont, S. and Hill, C.S. (2011) Recruitment of TIF1$γ$ to Chromatin via Its PHD Finger-Bromodomain Activates Its Ubiquitin Ligase and Transcriptional Repressor Activities. Mol Cell, 43, 85–96.

10. Champagne, K.S. and Kutateladze, T.G. (2009) Structural insight into histone recognition by the ING PHD fingers. Curr Drug Targets, 10, 432–441.

11. Jain, K., Fraser, C.S., Marunde, M.R., Parker, M.M., Sagum, C., Burg, J.M., Hall, N., Popova, I.K., Rodriguez, K.L., Vaidya, A., et al. (2020) Characterization of the plant homeodomain (PHD) reader family for their histone tail interactions. Epigenetics & Chromatin 2020 13:1, 13, 1–11.

12. Ettahar, A., Ferrigno, O., Zhang, M.-Z., Ohnishi, M., Ferrand, N., Prunier, C., Levy, L., Bourgeade, M.-F., Bieche, I., Romero, D.G., et al. (2013) Identification of PHRF1 as a Tumor Suppressor that Promotes the TGF-$β$ Cytostatic Program through Selective Release of TGIF-Driven PML Inactivation. Cell Rep, 4, 530–541.

13. Prunier, C., Zhang, M.-Z., Kumar, S., Levy, L., Ferrigno, O., Tzivion, G. and Atfi, A. (2015) Disruption of the PHRF1 Tumor Suppressor Network by PML-RAR$α$ Drives Acute Promyelocytic Leukemia Pathogenesis. Cell Rep, 10, 883–890.

14. Rebehmed, J., Revy, P., Faure, G., de Villartay, J.P. and Callebaut, I. (2014) Expanding the SRI domain family: a common scaffold for binding the phosphorylated C-terminal domain of RNA polymerase II. FEBS Lett, 588, 4431–4437.

15. Wang, Y., Wang, H., Pan, T., Li, L., Li, J., Yang, H., Wang, Y., Wang, H., Pan, T., Li, L., et al. (2016) Overexpression of PHRF1 attenuates the proliferation and tumorigenicity of non-small cell lung cancer cells. Oncotarget, 7, 64360–64370.

16. Harley, J.B., Alarcón-Riquelme, M.E., Criswell, L.A., Jacob, C.O., Kimberly, R.P., Moser, K.L., Tsao, B.P., Vyse, T.J., Langefeld, C.D., Langefeld, C.D., et al. (2008) Genome-wide association scan in women with systemic lupus erythematosus identifies susceptibility variants in ITGAM, PXK, KIAA1542 and other loci. Nat Genet, 40, 204–210.

17. Jarvinen, T.M., Hellquist, A., Zucchelli, M., Koskenmies, S., Panelius, J., Hasan, T., Julkunen, H., D’Amato, M. and Kere, J. (2012) Replication of GWAS-identified systemic lupus erythematosus susceptibility genes affirms B-cell receptor pathway signalling and strengthens the role of IRF5 in disease susceptibility in a Northern European population. Rheumatology, 51, 87–92.

18. Chang, C.-F., Chu, P.-C., Wu, P.-Y., Yu, M.-Y., Lee, J.-Y., Tsai, M.-D. and Chang, M.-S. (2015) PHRF1 promotes genome integrity by modulating non-homologous end-joining. Cell Death Dis, 6, e1716--e1716.

19. Jumper, J., Evans, R., Pritzel, A., Green, T., Figurnov, M., Ronneberger, O., Tunyasuvunakool, K., Bates, R., Žídek, A., Potapenko, A., et al. (2021) Highly accurate protein structure prediction with AlphaFold. Nature 2021 596:7873, 596, 583–589.

20. Rothbart, S.B., Krajewski, K., Nady, N., Tempel, W., Xue, S., Badeaux, A.I., Barsyte-Lovejoy, D., Martinez, J.Y., Bedford, M.T., Fuchs, S.M., et al. (2012) Association of UHRF1 with methylated H3K9 directs the maintenance of DNA methylation. Nat Struct Mol Biol, 19, 1155–1160.

21. Jain, K., Marunde, M.R., Burg, J.M., Gloor, S.L., Joseph, F.M., Gillespie, Z.B., Rodriguez, K.L., Howard, S.A., Popova, I.K., Hall, N.W., et al. (2022) An acetylation-mediated chromatin switch governs H3K4 methylation read-write capability. bioRxiv, 10.1101/2022.02.28.482307.

22. Morrison, E.A., Bowerman, S., Sylvers, K.L., Wereszczynski, J. and Musselman, C.A. (2018) The conformation of the histone H3 tail inhibits association of the BPTF PHD finger with the nucleosome. Elife, 7.

23. Lambert, J.P., Tucholska, M., Go, C., Knight, J.D.R. and Gingras, A.C. (2015) Proximity biotinylation and affinity purification are complementary approaches for the interactome mapping of chromatin-associated protein complexes. J Proteomics, 118, 81–94.

24. Rappsilber, J., Mann, M. and Ishihama, Y. (2007) Protocol for micro-purification, enrichment, pre-fractionation and storage of peptides for proteomics using StageTips. Nature Protocols 2007 2:8, 2, 1896–1906.

25. Shah, A.D., Goode, R.J.A., Huang, C., Powell, D.R. and Schittenhelm, R.B. (2019) Lfq-Analyst: An easy-To-use interactive web platform to analyze and visualize label-free proteomics data preprocessed with maxquant. J Proteome Res, 10.1021/ACS.JPROTEOME.9B00496/SUPPL_FILE/PR9B00496_SI_001.PDF.

26. Cox, J., Hein, M.Y., Luber, C.A., Paron, I., Nagaraj, N. and Mann, M. (2014) Accurate proteome-wide label-free quantification by delayed normalization and maximal peptide ratio extraction, termed MaxLFQ. Molecular and Cellular Proteomics, 13, 2513–2526.

27. Dobin, A., Davis, C.A., Schlesinger, F., Drenkow, J., Zaleski, C., Jha, S., Batut, P., Chaisson, M. and Gingeras, T.R. (2013) STAR: ultrafast universal RNA-seq aligner. Bioinformatics, 29, 15–21.

28. Love, M.I., Huber, W. and Anders, S. (2014) Moderated estimation of fold change and dispersion for RNA-seq data with DESeq2. Genome Biol, 15, 1–21.

29. Shen, S., Park, J.W., Lu, Z.X., Lin, L., Henry, M.D., Wu, Y.N., Zhou, Q. and Xing, Y. (2014) rMATS: Robust and flexible detection of differential alternative splicing from replicate RNA-Seq data. Proc Natl Acad Sci U S A, 111, E5593–E5601.

30. Huang, D.W., Sherman, B.T. and Lempicki, R.A. (2008) Systematic and integrative analysis of large gene lists using DAVID bioinformatics resources. Nature Protocols 2009 4:1, 4, 44–57.

31. Kuleshov, M. V., Jones, M.R., Rouillard, A.D., Fernandez, N.F., Duan, Q., Wang, Z., Koplev, S., Jenkins, S.L., Jagodnik, K.M., Lachmann, A., et al. (2016) Enrichr: a comprehensive gene set enrichment analysis web server 2016 update. Nucleic Acids Res, 44, W90–W97.

32. Tsherniak, A., Vazquez, F., Montgomery, P.G., Weir, B.A., Kryukov, G., Cowley, G.S., Gill, S., Harrington, W.F., Pantel, S., Krill-Burger, J.M., et al. (2017) Defining a Cancer Dependency Map. Cell, 170, 564–576.e16.

33. DepMap 24Q2 Public.

34. Kandoth, C., McLellan, M.D., Vandin, F., Ye, K., Niu, B., Lu, C., Xie, M., Zhang, Q., McMichael, J.F., Wyczalkowski, M.A., et al. (2013) Mutational landscape and significance across 12 major cancer types. Nature 2013 502:7471, 502, 333–339.

35. Aguet, F., Brown, A.A., Castel, S.E., Davis, J.R., He, Y., Jo, B., Mohammadi, P., Park, Y.S., Parsana, P., Segrè, A. V., et al. (2017) Genetic effects on gene expression across human tissues. Nature 2017 550:7675, 550, 204–213.

36. McQuin, C., Goodman, A., Chernyshev, V., Kamentsky, L., Cimini, B.A., Karhohs, K.W., Doan, M., Ding, L., Rafelski, S.M., Thirstrup, D., et al. (2018) CellProfiler 3.0: Next-generation image processing for biology. PLoS Biol, 16, e2005970.

37. Fasiolo, M., Wood, S.N., Zaffran, M., Nedellec, R. and Goude, Y. (2021) Fast Calibrated Additive Quantile Regression. J Am Stat Assoc, 116, 1402–1412.

38. Boamah, D., Lin, T., Poppinga, F.A., Basu, S., Rahman, S., Essel, F. and Chakravarty, S. (2018) Characteristics of a PHD Finger Subtype. Biochemistry, 57, 525–539.

39. Tate, J.G., Bamford, S., Jubb, H.C., Sondka, Z., Beare, D.M., Bindal, N., Boutselakis, H., Cole, C.G., Creatore, C., Dawson, E., et al. (2019) COSMIC: the Catalogue Of Somatic Mutations In Cancer. Nucleic Acids Res, 47, D941–D947.

40. Weinberg, D.N., Papillon-Cavanagh, S., Chen, H., Yue, Y., Chen, X., Rajagopalan, K.N., Horth, C., McGuire, J.T., Xu, X., Nikbakht, H., et al. (2019) The histone mark H3K36me2 recruits DNMT3A and shapes the intergenic DNA methylation landscape. Nature, 573, 281–286.

41. Marunde, M.R., Popova, I.K., Weinzapfel, E.N. and Keogh, M.-C. (2022) The dCypher Approach to Interrogate Chromatin Reader Activity Against Posttranslational Modification-Defined Histone Peptides and Nucleosomes. 10.1007/978-1-0716-2140-0_13.

42. Marunde, M.R., Fuchs, H.A., Burg, J.M., Popova, I.K., Vaidya, A., Hall, N.W., Weinzapfel, E.N., Meiners, M.J., Watson, R., Gillespie, Z.B., et al. (2024) Nucleosome conformation dictates the histone code. Elife, 13.

43. Jain, K., Marunde, M.R., Burg, J.M., Gloor, S.L., Joseph, F.M., Poncha, K.F., Gillespie, Z.B., Rodriguez, K.L., Popova, I.K., Hall, N.W., et al. (2023) An acetylation-mediated chromatin switch governs H3K4 methylation read-write capability. Elife, 12.

44. Abuetabh, Y., Wu, H.H., Chai, C., Al Yousef, H., Persad, S., Sergi, C.M. and Leng, R. (2022) DNA damage response revisited: the p53 family and its regulators provide endless cancer therapy opportunities. Experimental & Molecular Medicine 2022, 10.1038/s12276-022-00863-4.

45. Sharp, P.A., Chakraborty, A.K., Henninger, J.E. and Young, R.A. (2022) RNA in formation and regulation of transcriptional condensates. RNA, 28, 52–57.

46. Kizer, K.O., Phatnani, H.P., Shibata, Y., Hall, H., Greenleaf, A.L. and Strahl, B.D. (2005) A Novel Domain in Set2 Mediates RNA Polymerase II Interaction and Couples Histone H3 K36 Methylation with Transcript Elongation. Mol Cell Biol, 25, 3305–3316.

47. Luco, R.F., Pan, Q., Tominaga, K., Blencowe, B.J., Pereira-Smith, O.M. and Misteli, T. (2010) Regulation of alternative splicing by histone modifications. Science (1979), 327, 996–1000.

48. Krämer, A., Green, J., Pollard, J. and Tugendreich, S. (2014) Causal analysis approaches in Ingenuity Pathway Analysis. Bioinformatics, 30, 523–530.

49. Chankova, S.G., Dimova, E., Dimitrova, M. and Bryant, P.E. (2007) Induction of DNA double-strand breaks by zeocin in Chlamydomonas reinhardtii and the role of increased DNA double-strand breaks rejoining in the formation of an adaptive response. Radiat Environ Biophys, 46, 409–416.

50. Povirk, L.F. (1996) DNA damage and mutagenesis by radiomimetic DNA-cleaving agents: bleomycin, neocarzinostatin and other enediynes. Mutation Research/Fundamental and Molecular Mechanisms of Mutagenesis, 355, 71–89.

51. Alabert, C., Bianco, J.N. and Pasero, P. (2009) Differential regulation of homologous recombination at DNA breaks and replication forks by the Mrc1 branch of the S-phase checkpoint. EMBO Journal, 28, 1131–1141.

52. Huang, R.X. and Zhou, P.K. (2020) DNA damage response signaling pathways and targets for radiotherapy sensitization in cancer. Signal Transduction and Targeted Therapy 2020 5:1, 5, 1–27.

53. Nikfarjam, S. and Singh, K.K. (2022) DNA damage response signaling: A common link between cancer and cardiovascular diseases. Cancer Med, 10.1002/CAM4.5274.

54. Jackson, S.P. and Bartek, J. (2009) The DNA-damage response in human biology and disease. Nature 2009 461:7267, 461, 1071–1078.

55. Ahmed, D., Eide, P.W., Eilertsen, I.A., Danielsen, S.A., Eknæs, M., Hektoen, M., Lind, G.E. and Lothe, R.A. (2013) Epigenetic and genetic features of 24 colon cancer cell lines. Oncogenesis 2013 2:9, 2, e71–e71.

56. Lin, H.-W., Shih, T.-W., Amanna, A. And Chang, M.-S. (2023) PHRF1 Promotes Cell Invasion by Modulating SOX4 Expression in Colorectal Cancer HCT116-p53−/− Cells. Anticancer Res, 43, 5437–5446.

57. Lee, J.Y., Fan, C.C., Chou, N.L., Lin, H.W. and Chang, M.S. (2020) PHRF1 promotes migration and invasion by modulating ZEB1 expression. PLoS One, 15, e0236876.

58. David, C.J. and Manley, J.L. (2010) Alternative pre-mRNA splicing regulation in cancer: pathways and programs unhinged. Genes Dev, 24, 2343–2364.

59. Oltean, S. and Bates, D.O. (2013) Hallmarks of alternative splicing in cancer. Oncogene 2014 33:46, 33, 5311–5318.

60. Anczukow, O. and Krainer, A.R. (2016) Splicing-factor alterations in cancers. RNA, 22, 1285–1301.

61. Dvinge, H., Kim, E., Abdel-Wahab, O. and Bradley, R.K. (2016) RNA splicing factors as oncoproteins and tumour suppressors. Nature Reviews Cancer 2016 16:7, 16, 413–430.

62. Ciccia, A. and Elledge, S.J. (2010) The DNA Damage Response: Making It Safe to Play with Knives. Mol Cell, 40, 179–204.

