## Supplementary Material for "Histone H3 N-terminal recognition by the PHD finger of PHRF1 is required for proper DNA damage response"

**SUPPLEMENTARY MATERIAL – Jain et al. 2024.**

Supplementary File 1. Binding data from the histone peptide microarray.

Supplementary File 2. BioID MS results.

Supplementary File 3. BioID DAVID GO terms.

Supplementary File 4. HeLa ΔPHRF1 v. control RNAseq differentially expressed genes.

Supplementary File 5. HeLa RNAseq Ingenuity Pathway Analysis.

Supplementary File 6. Summary of rMATS data.

Supplementary File 7. Enrichr GO terms from rMATS data.

Supplementary File 8. HCT116 ΔPHRF1 v. control RNAseq differentially expressed genes.

Supplementary File 9. HCT116 RNAseq Ingenuity Pathway Analysis.

**Supplemental Table S1. Statistical parameters of qGAM analysis for γH2A.X intensity/nucleus in HeLa cells**

| Comparison | Parametric Coefficient | p-value | LRT p-value |
|---|---|---|---|
| ΔPHRF1 vs Control | -345.25 | $4.79 \times 10^{-5}$ (****) | $3.26 \times 10^{-10}$ |

Key:

- **Parametric Coefficient:** Difference in overall baseline γH2A.X intensity between cell lines.
- **p-value**: Statistical significance of Parametric Coefficient determined by a Wald Test.
- **LRT:** Likelihood Ratio test, used to assess differences in patterns between cell lines.

**Supplemental Table S2. Statistical parameters of qGAM analysis for γH2A.X and 53BP1 foci/nucleus HCT116 cells**

| | Comparison | Parametric Coefficient | p-value | LRT p-value |
|---|---|---|---|---|
| **γH2A.X foci/nucleus** | ΔPHRF1 vs Control | -1.9417 | $< 2.2 \times 10^{-16}$ (****) | $4.78 \times 10^{-9}$ |
| **53BP1 foci/nucleus** | ΔPHRF1 vs Control | -3.4801 | $< 2.2 \times 10^{-16}$ (****) | $< 2.2 \times 10^{-16}$ |

Key:

- **Parametric Coefficient:** Difference in overall baseline foci counts between cell lines.
- **p-value**: Statistical significance of Parametric Coefficient determined by a Wald Test.

- **LRT:** Likelihood Ratio test, used to assess differences in patterns between cell lines.

**Supplemental Table S3. Statistical parameters of qGAM analysis for γH2A.X and 53BP1 foci/nucleus HCT116 complimented cell lines**

| | **Comparison** | **Parametric Coefficient** | **p-value** | **LRT p-value** |
|---|---|---|---|---|
| **γH2A.X foci/nucleus** | ΔPHRF1+EV vs Control+EV | -5.405 | 0.00115 (***) | $< 2.2 \times 10^{-16}$ |
| | ΔPHRF1+EV vs ΔPHRF1+3XFlag-PHRF1$^{WT}$ | -7.175 | $1.87 \times 10^{-7}$ (****) | $< 2.2 \times 10^{-16}$ |
| | ΔPHRF1+EV vs ΔPHRF1+3XFlag-PHRF1$^{P221L}$ | -0.787 | 0.584 (ns) | $< 2.2 \times 10^{-16}$ |
| **53BP1 foci/nucleus** | ΔPHRF1+EV vs Control+EV | -3.256 | $2.37 \times 10^{-11}$ (****) | $2.41 \times 10^{-10}$ |
| | ΔPHRF1+EV vs ΔPHRF1+3XFlag-PHRF1$^{WT}$ | -2.425 | $1.89 \times 10^{-10}$ (****) | $8.71 \times 10^{-6}$ |
| | ΔPHRF1+EV vs ΔPHRF1+3XFlag-PHRF1$^{P221L}$ | 0.44 | 0.262 (ns) | $2.49 \times 10^{-10}$ |

Key:

- **Parametric Coefficient:** Difference in overall baseline foci counts between cell lines.
- **p-value**: Statistical significance of Parametric Coefficient determined by a Wald Test.
- **LRT:** Likelihood Ratio test, used to assess differences in patterns between cell lines.
- **ns:** not significant

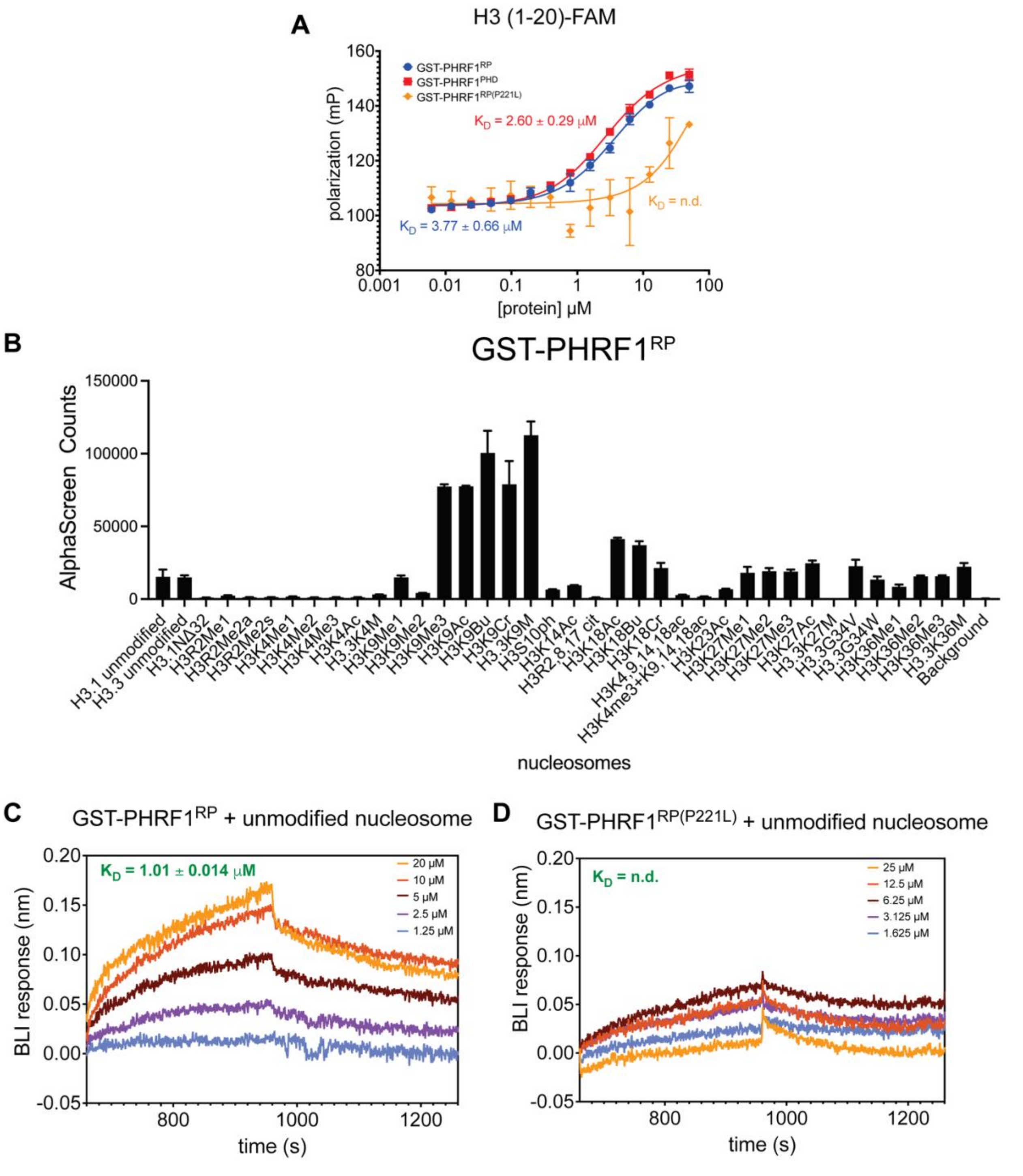


**Supplementary Figure S1. Binding preference of PHRF1 with histone H3 peptides and nucleosomes. A.** Fluorescence polarization binding assays of H3 (1-20) (C-terminally 5-FAM-labeled) with GST-PHRF1$^{RP}$ (red), GST-PHRF1$^{PHD}$ (blue), and GST-PHRF1$^{RP(P221L)}$ (orange). $K_D$ values are indicated where calculable, error bars represent S.D., n = 3, and n.d. = not determined. **B.** dCypher Alphascreen counts for the interaction of GST-PHRF1$^{RP}$ (3 nM) with PTM-defined biotinylated nucleosomes. All error bars represent the range of two replicates. Key: H3.1 NΔ2 and NΔ32 are nucleosomes lacking residues 1-2 or 1-32 of H3, respectively. **C.** Biolayer interferometry (BLI) response for the interaction between GST-PHRF1$^{RP}$ and H3 unmodified nucleosome (12.5 nM). **D.** BLI response for the interaction between GST-PHRF1$^{P221L}$ and H3 unmodified nucleosome (12.5 nM). For both **C** and **D**, the time window representing the association and dissociation phases for each curve are color coded by protein concentration. $K_D$ values are determined from a global fit using the calculated $k_{on}$ and $k_{off}$ rates for each experiment; n.d. = not determined.

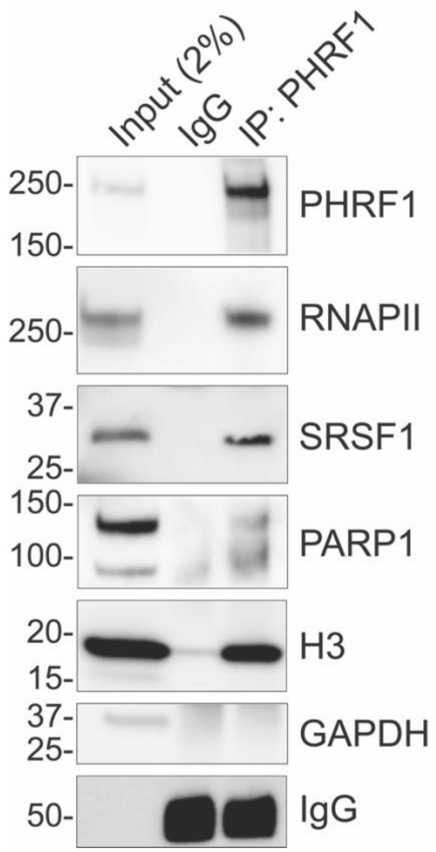


**Supplementary Figure S2. PHRF1 associates with splicing, transcription, and DDR-related proteins in cells.** Western blot analysis of splicing, transcription, and DDR-associated proteins following immunoprecipitation of endogenous PHRF1 in HeLa cells with IgG as a negative control.

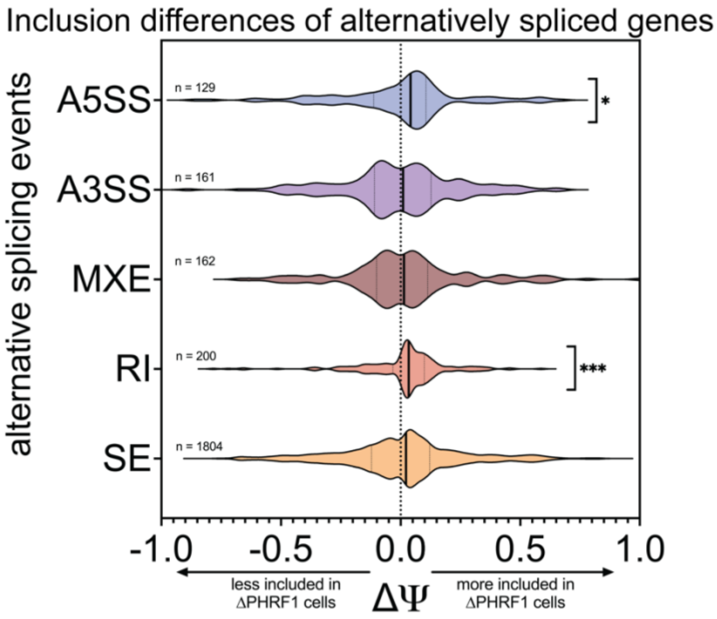


**Supplementary Figure S3. Significant increase in inclusion of A5SS and RI splicing events upon loss of PHRF1.** Comparison of alternative splicing events in ΔPHRF1 cells versus control cells relative to no change ($\Delta\Psi = 0$). Statistical significance was determined by the Wilcoxon signed-rank test; * = p-value < 0.05, *** = p-value < 0.001. Key: SE: Skipped Exons, RI: Retained Introns, MXE: Mutually Exclusive Exons, A3SS: Alternative 3’ Splice Sites, A5SS: Alternative 5’ Splice Sites, and $\Delta\Psi$: represents the difference in Percent Spliced In (PSI) between the two conditions.

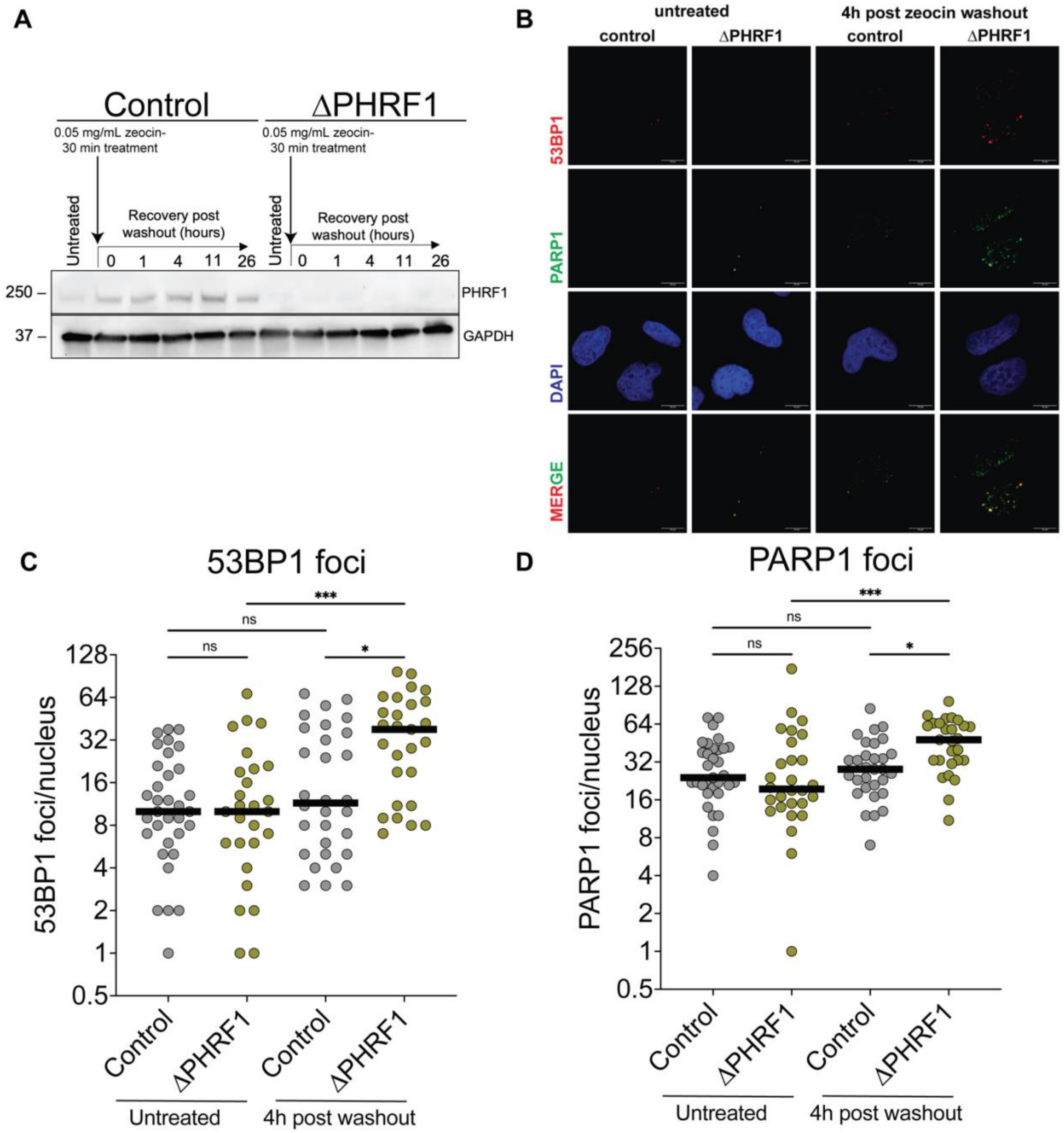


**Supplementary Figure S4. PHRF1 affects in cellular DDR. A.** Representative western blot showing PHRF1 protein levels over time during the zeocin DDR assay described in **Figure 4A. B.** Representative confocal immunofluorescence of control and ΔPHRF1 cells stained with anti-53BP1 (red), anti-PARP1 (green) antibodies, and DAPI (blue) at indicated time points along the zeocin DDR assay. The scale bar is 10 μm. **C, D.** Quantitative analysis of 53BP1(**C**) and PARP1 (**D**) foci/nuclear in control (gray) and ΔPHRF1 (mustard green) cells from a zeocin DDR assay.

Medians are shown as solid black lines and compared using a Kruskal-Wallis test with a Dunn's test for multiple comparisons. *** = p-value < 0.0002, * = p-value < 0.05, ns = not significant.

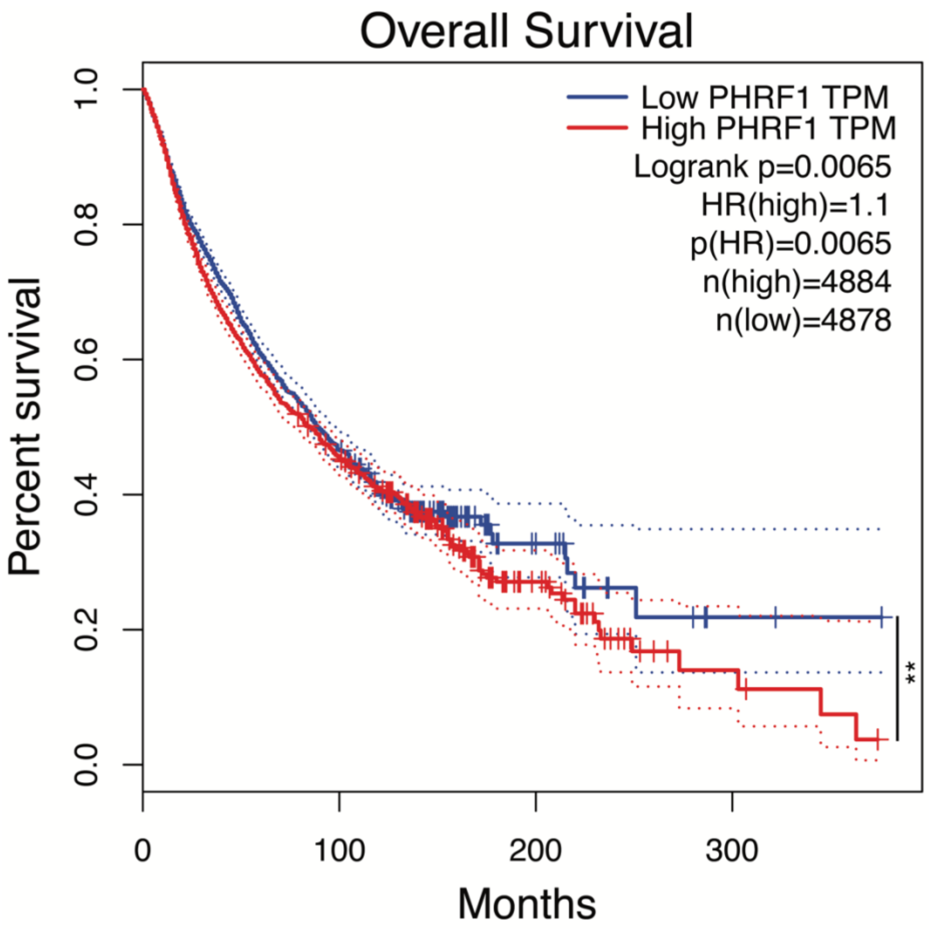


**Supplementary Figure S5. Correlation of PHRF1 expression levels on overall cancer survival in patients.** Kaplan-Meier survival curves comparing patients from The Cancer Genome Atlas (TCGA) with high (red solid line) and low (blue solid line) expression of PHRF1, with 95% confidence intervals indicated by dotted lines. Statistical parameters for survival are indicated on the graph. ** = p-value < 0.01.

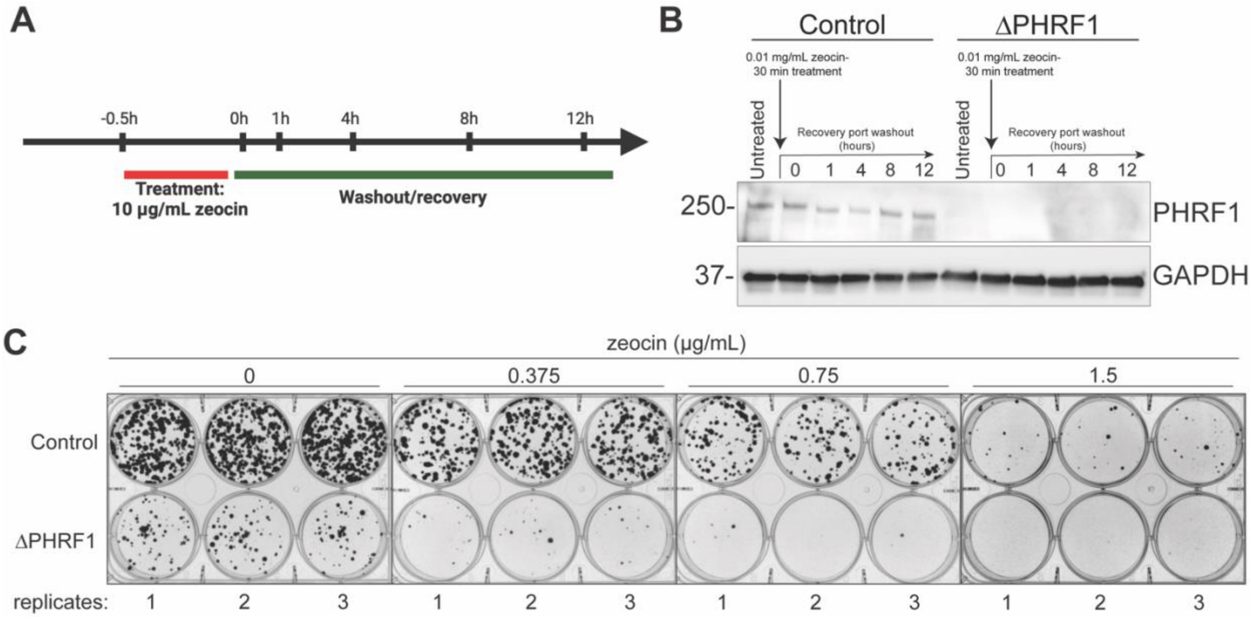


**Supplementary Figure S6. Zeocin DDR assay in HCT116 control and ∆PHRF1 cells. A.** Experimental schematic of zeocin DDR assay in HCT116 cells. **B.** Representative western blot of PHRF1 protein levels in control and ∆PHRF1 HCT116 cells during a zeocin DDR assay. **C.** All images of the CFA experiment shown in **Figure 6F and 6G**. replicate numbers are shown at the bottom of each image.

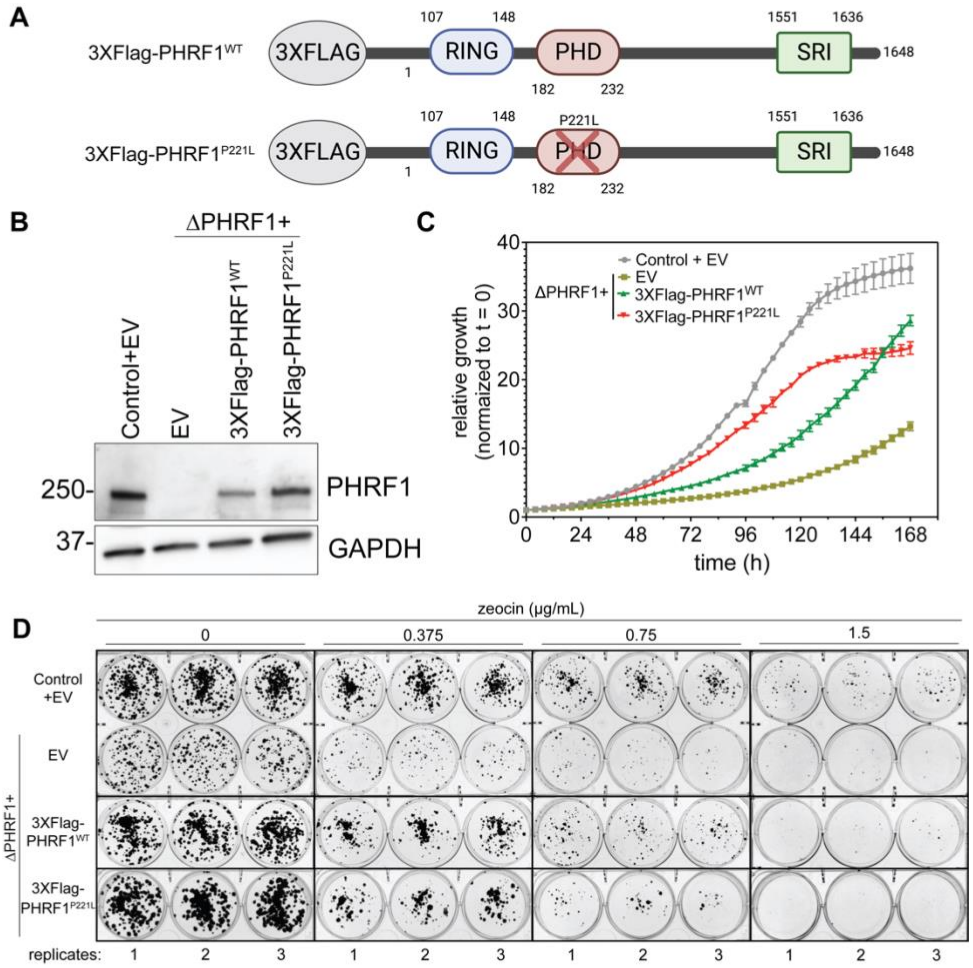


**Supplementary Figure S7. PHRF1 complementation analyses of cellular growth in HCT116.** **A.** Sequence schematic showing the PHRF1 constructs that were incorporated into ∆PHRF1 HCT116 through lentiviral infection. **B.** Representative western blot showing PHRF1 protein levels in four HCT116 cell lines: Control+EV, ∆PHRF1+EV, ∆PHRF1+3XFlag-PHRF1$^{WT}$, and ∆PHRF1+3XFlag-PHRF1$^{P221L}$. **C.** Growth curves measured through live-cell imaging on the Incucyte S3 of the four cell lines from panel **B**. Error bars represent S.D. and n = 3. **D.** All images of the CFA experiment shown in **Figure 7A and 7B**. Replicate numbers are shown at the bottom of each image.

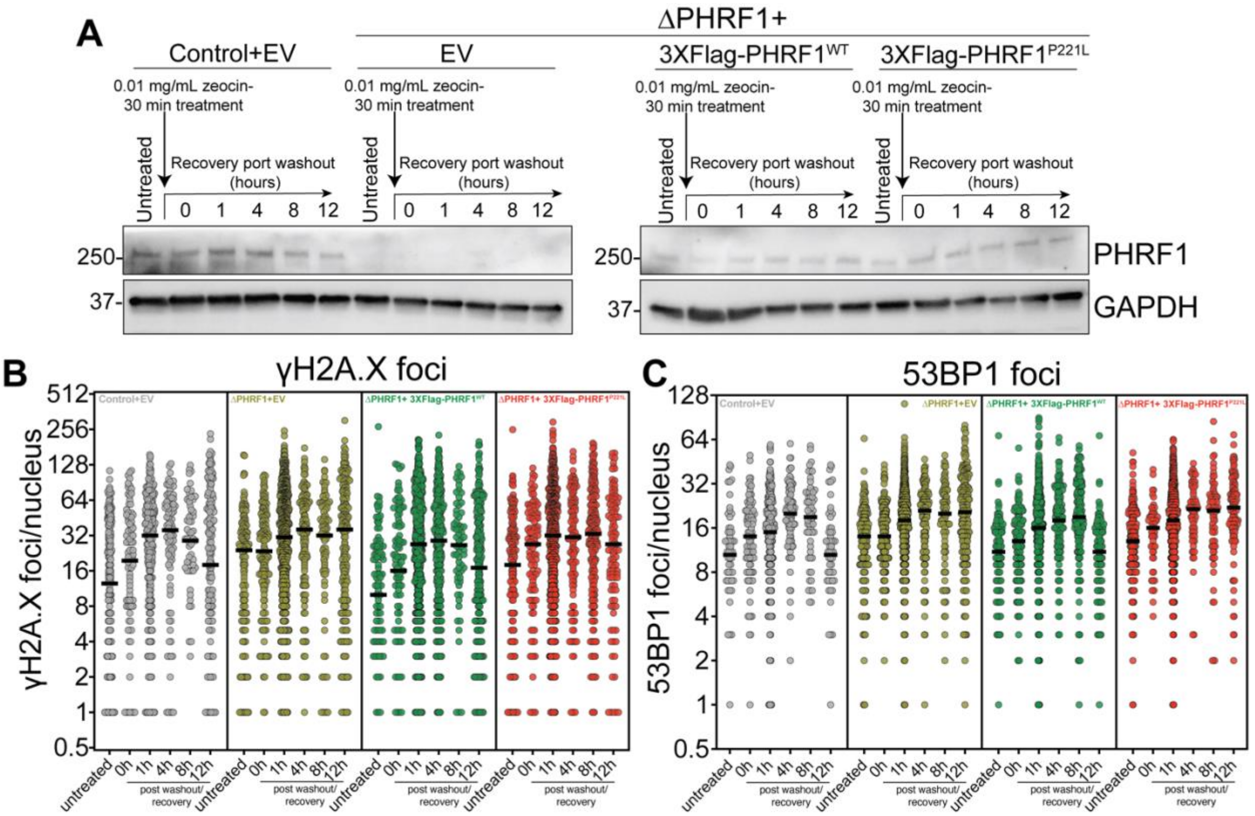


**Supplementary Figure S8. PHRF1 complementation analyses of cellular DDR in HCT116. A.** Representative western blot of PHRF1 protein levels in the indicated HCT116 cell lines during a zeocin DDR assay. **B, C.** Quantitative analysis of γH2A.X (**B**) and 53BP1 (**C**) foci/ nucleus in Control+EV (gray), ∆PHRF1+EV (mustard green), ∆PHRF1+3XFlag-PHRF1$^{WT}$ (green), and ∆PHRF1+3XFlag-PHRF1$^{P221L}$ (red) HCT116 cells. Median values are shown with solid black lines.
